# The elongation of *Mest* transcript into *MestXL* sustains, but does not initiate, the maternal allele bias of its convergent gene *Copg2* during neurogenesis

**DOI:** 10.64898/2026.03.13.711300

**Authors:** Sophie Perillous, Anne-Charlotte Fromaget, Céline Gonthier-Guéret, Ornella Clerici, Marion Espenel, Alice Murigneux, Sophie Phan, Lea Feit, Catherine Vaurs-Barriére, Davide Normanno, Amanda Ha, Nolan Ashworth, Aaron B. Bogutz, Hiromi Kamura, Nuttamonpat Gumpangseth, Kenichiro Hata, Bertille Montibus, Kazuhiko Nakabayashi, Louis Lefebvre, Tristan Bouschet, Franck Court, Philippe Arnaud

**Author notes:** Corresponding authors (PA), (FC); (TB). These authors contribute equally (Sophie Perillous, Anne-Charlotte Fromaget).

## Abstract

Precise gene dosage control is critical for establishing cellular identity and development, especially for imprinted genes, where dosage imbalances are linked to severe pathologies such as neurodevelopmental disorders. The *Mest/Copg2* imprinted locus is a paradigm for this fine-tuned regulation. While *Mest* is constitutively expressed from the paternal allele, *Copg2* expression shifts from biallelic to a maternal allele bias specifically during neural differentiation, a transition proposed to involve transcriptional interference mediated by the long *Mest* isoform, *MestXL*, which extends into the *Copg2* locus. However, the mechanisms underlying this allelic switch, and whether factors beyond *MestXL* contribute, are elusive. To address this, we employed a stem cell-based brain organoid model, integrating multi-omic analyses, 3D chromatin structure mapping, and functional approaches to dissect the regulatory events governing the induction and maintenance of *Copg2* maternal allele bias throughout neural lineage specification.

Our findings challenge the prevailing model by demonstrating that the maternal allele bias of *Copg2* during neural differentiation is not solely driven by *MestXL*-mediated transcriptional interference. Instead, our data support a temporal and neural stage-specific two-step mechanism: putative enhancer-driven activation of the maternal allele in neural progenitor cells is followed by *MestXL*-dependent repression of the paternal allele in neuron-enriched stages. This uncovers an unexpected layer of complexity in the regulation of imprinted gene dosage during brain development, with profound implications for understanding the molecular underpinnings of neurodevelopmental disorders.

## Introduction

Establishing cellular identity during development relies on the precise integration of developmental cues and environmental signals into tightly regulated gene expression programs^1^. Central to this process is the accurate control of gene dosage, where even minor disruptions can lead to severe human pathologies, including cancer, autoimmune diseases, and neurological disorders^2^. Among the most dosage-sensitive genes are imprinted genes, whose quantitative regulation is essential for normal development and physiological function.

Genomic imprinting is an epigenetic mechanism that restricts the expression of certain mammalian genes to a single parental allele. Of the approximately 200 imprinted genes identified, many play pivotal roles in cell proliferation, fetal and placental growth, energy homeostasis, and metabolic adaptation^3^. Notably, imprinted genes are also critical for brain function and behaviour, with their dysregulation linked to neurobehavioral disorders such as Prader-Willi and Angelman syndromes^4^. While traditionally viewed as strictly monoallelic, advances in high-resolution RNA-seq and single-cell technologies have revealed a more complex landscape: many imprinted genes exhibit parent-of-origin allelic bias rather than complete monoallelic expression^5,6^. The functional significance of this quantitative regulation is exemplified by the paternally biased gene *Bcl2l1*, where brain-specific deletion of the paternal, but not maternal, allele results in the loss of specific neuronal subtypes, highlighting the importance of precise allelic dosage in development and adult physiology^5^.

In both humans and mice, imprinted genes are often organized into evolutionarily conserved genomic clusters spanning several megabases. These genomic domains contain multiple imprinted genes whose allele-specific expression is primarily regulated by a key imprinting control region (ICRs). Each ICR overlaps with a gametic differentially methylated region (gDMR) that acquires parent-specific methylation in the germline and is maintained throughout development. This constitutive methylation orchestrates monoallelic or biased expression through a combination of direct and long-range regulatory mechanisms, many of which are tissue- or stage-specific, resulting in complex spatiotemporal expression patterns^7^.

ICRs can regulate distal gene expression by acting as allele-specific insulators, modulating long-range enhancer–promoter interactions within imprinted domains. These interactions are structured within topologically associating domains (TADs). For example, at the *H19* and *Peg13* loci, binding of the methylation-sensitive insulator protein CTCF to the unmethylated ICR allele establishes allele-specific sub-TAD configurations, contributing to parent-of-origin transcriptional programs^8–10^. The repeated observation of CTCF enrichment at distinct ICRs^11^, suggests that allele-specific chromatin organization may be a conserved strategy for coordinating imprinted gene expression across large genomic regions.

Beyond insulator function, ICRs also exert regulatory effects through their promoter activity and transcriptional outputs^7^. At the *Igf2r* and *Kcnq1* loci, paternally expressed long non-coding RNAs (lncRNAs) *Airn* and *Kcnq1ot1*, respectively, mediate long-range cis-repression of neighbouring genes over distances of up to 10 Mb and 800 kb, respectively^6^. Additionally, allele-specific transcription initiated at ICRs can interfere with overlapping genes, as seen for instance at loci like *Igf2r*, *Nap1l5* and *Snrpn*, where transcriptional overlap modulates expression through mechanisms such as promoter occlusion^12^, alternative polyadenylation^13,14^, or transcriptional collision^15,16^.

A compelling example of a postulated transcriptional interference regulation is found within the *Mest/Copg2* imprinted domain. Located on mouse chromosome 6, syntenic to human chromosome 7, this approximately 250 kb domain encompasses several imprinted genes, including the paternally expressed gene *Mest* (*Mesoderm-specific transcript*, also known as *Peg1*) and the maternally biased gene *Copg2*^16^. Regulation of this domain is mediated by a maternally methylated imprinting control region at the *Mest* promoter. *Mest* encodes an alpha/beta hydrolase involved in lipid metabolism and embryonic development^17,18^, while *Copg2* (*Coatomer Protein Complex Subunit Gamma 2*) encodes a subunit of the COPI coatomer complex, which is essential for retrograde vesicular trafficking and Golgi structure maintenance^19^. In humans, *MEST* has been linked to cognitive ability^20^, autism susceptibility^21^, and neurodevelopmental disorders, including reported *de novo* paternal 7q32.2 microdeletion in patients with Silver–Russell syndrome^22,23^. Consistent with these findings, paternal deletion of *Mest* in mice results in growth retardation^18,24^. Additionally, genetic alterations affecting *COPG2* have been associated with neurodevelopmental disorders, such as intellectual disability and autism spectrum disorders^25^. Collectively, these observations underscore the critical role of this domain in developmental and neurodevelopmental processes.

Convergent transcription at the *Mest*/*Copg2* locus further illustrates the complexity of imprinted gene regulation. During neural differentiation, *Copg2* expression shifts from biallelic to a maternal allele bias^16^, a transition proposed to involve transcriptional interference mediated by a long *Mest* isoform, *MestXL*, which extends far into the *Copg2* locus. However, the precise mechanisms underlying this switch in *Copg2* allelic usage, and whether factors beyond *MestXL* contribute to this process, remain unresolved.

To address these questions, we leveraged a brain organoid system in which the induction of *MestXL* and the maternal allele bias of *Copg2* are recapitulated. Our findings redefine the regulatory framework governing *Copg2*’s maternal allele bias during neural differentiation. We demonstrate that the initiation of the allelic switch occurs independently of *MestXL*, and reveal that temporal and neural stage-specific regulatory mechanisms collaborate to fine-tune *Copg2* allelic expression throughout neural lineage formation.

## Results

### Mouse brain organoids as a model for studying *MestXL* induction and allelic usage at *Copg2* during neural development

To investigate the regulatory mechanisms governing the interaction between *Mest* and *Copg2* during neural development, we utilized mouse brain organoids as a model system. This model, which we recently characterized, faithfully recapitulates the *in vivo* gene expression profiles observed in the newborn mice brain^26^. Hybrid embryonic stem cells (mESCs) derived from reciprocal crosses between C57BL/6J (B6) and JF1 mice (designated as B/J) were differentiated into brain organoids over 21 days (Fig. 1A). As shown by immunofluorescence (Fig.1A) and RT-qPCR (Suppl Fig1A) experiments, this differentiation paradigm led to the sequential emergence of distinct cellular populations: neural progenitors expressing *Nestin* and *Pax6* at day 7, followed at day 14 by radial glia (FABP7+) and neurons (expressing *Tubb3* or *Elavl3*), and, at day 21, by neurons (including TBR1+ cells) and astrocytes (expressing *Gfap*) (Fig.1A and Suppl Fig1A).

**Figure 1.**
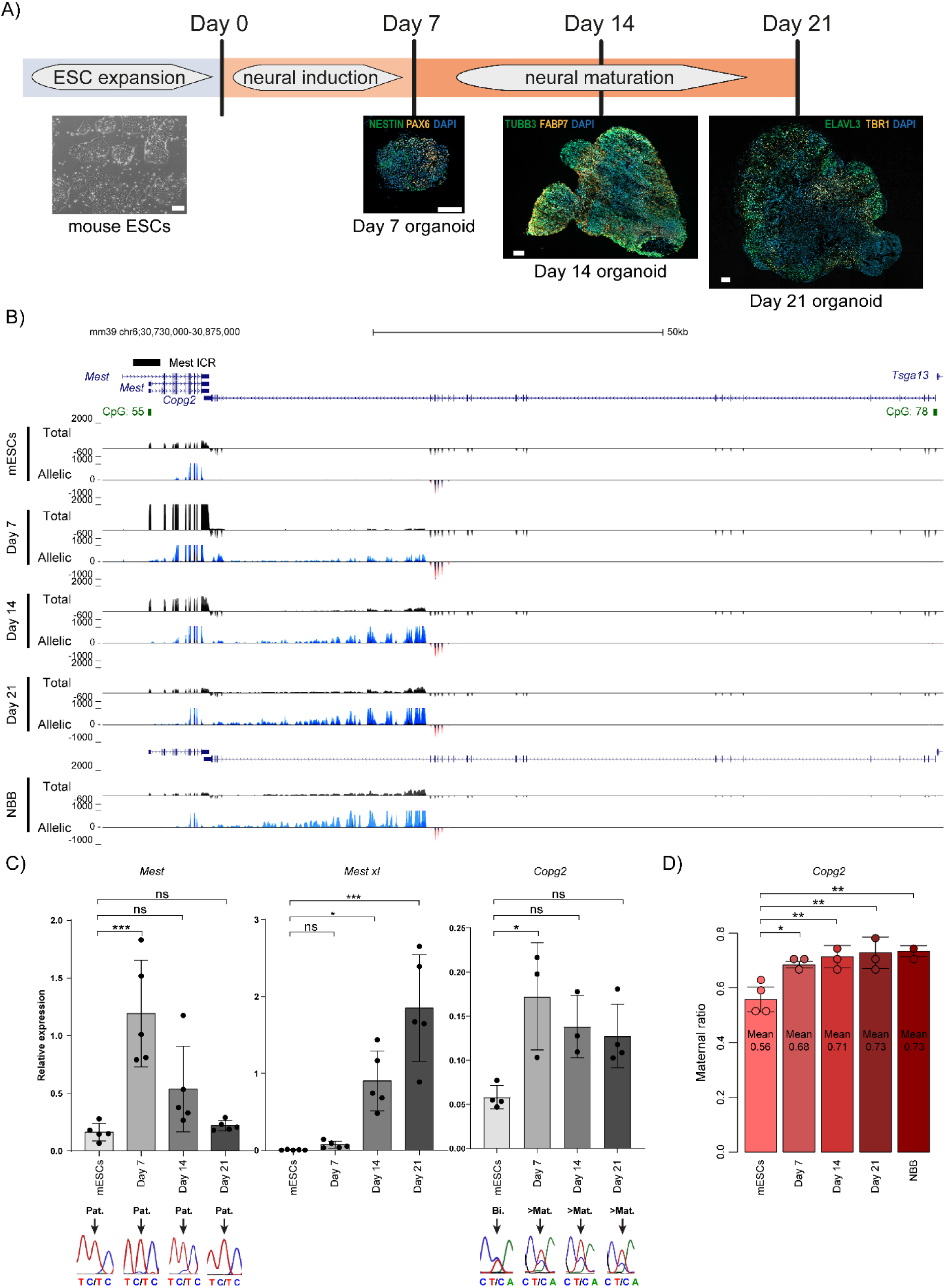
Expression patterns within the *Mest* domain in brain organoids recapitulate in vivo patterns. **A)** Schematic timeline of brain organoids generation from mESCs. Immunostaining of mESCs-derived brain organoids : cryosections were stained with NESTIN and PAX6 (at Day 7), TUBB3 and FABP7 (at day 14) and ELAVL3 and TBR1 (at Day 21). Scale bars: 10 µm for mESCs, and 100 µm for all cryosections on organoids. **B)** Genome browser view at the *Mest* and *Copg2* genes to show the allelic-oriented RNA-seq signals in B/J ESCs (n=4) and at day 7 (*n* = 3), day 14 (*n* = 3), and day 21 (*n* = 3) of in vitro brain organoid differentiation, as well as in merged B/J and J/B newborn brain tissue (NBB) (*n* = 2). For each condition, the quantitative and merged parental allelic RNA-seq signals are at the top and bottom, respectively. Maternal and paternal expression levels are shown in red and blue, respectively. **C)** RT-qPCR analysis of *Mest*, *MestXL*, and *Copg2* expression levels in B/J ESCs and at Day 7, Day 14 and Day 21 of in vitro brain organoid differentiation. Parental origin of expression is indicated for *Mest* and *Copg2*. Values are the mean of three to five independent experiments, each analyzed in duplicate. Data are presented as a percentage of expression relative to the geometric mean of three housekeeping genes (*Gapdh*, *Gus*, and *Tbp*), and are shown as mean ± SEM. Parental origin was determined by direct sequencing of sample-specific PCR products, leveraging strain-specific SNPs in the analyzed regions. **D)** Ratio of maternal expression at the *Copg2* locus, calculated as the percentage of RNA-seq reads derived from the maternal allele in B/J ESCs (*n* = 4), day 7 (*n* = 3), day 14 (*n* = 3), day 21 (*n* = 3) of in vitro brain organoid differentiation, and in newborn brain tissue (NBB) (*n* = 2). In C) and D), Statistical significance was determined with the one-way ANOVA and a post-hoc Tukey test (one-way ANOVA p<0.05 and Tukey test p < 0.05 *, p < 0.01 **, p < 0.001 ***).

RNA-seq and RT-qPCR analyses at the *Mest/Copg2* domain revealed that, in ESCs, imprinted expression was restricted to *Mest*, which exhibited exclusively paternal expression, while *Copg2* was expressed biallelically. During differentiation, *Mest* maintained its paternal expression but displayed dynamic changes: a significant increase at day 7, followed by a decline at days 14 and 21. Concurrently, its long isoform, *MestXL*, which we found also paternally expressed, was detected from day 7, coinciding with the peak expression of *Mest*, and gradually increased up to day 21 (Fig. 1B & C).

This *MestXL* transcript overlaps the *Copg2* transcriptional domain, spanning nearly 40 kb and extending to the region downstream of exon 20 (out of 24) in *Copg2* (Fig. 1B). Consistent with this observation, in the newborn brain, a strong paternal signal for H3K36me3, a histone mark associated with transcriptional elongation, was detected spanning from *Mest* across a ∼40 kb region within *Copg2*. In contrast, the region of *Copg2* from exon 1 to 20 exhibited biallelic enrichment for H3K36me3 (Supp. Fig. 1B).

In parallel with the peak of expression of *Mest* and the emergence of *MestXL*, *Copg2* shifted from a biallelic to a preferentially maternal expression, accompanied by an overall increase in Copg2 transcript levels at day 7, followed by a relative decline at day 14 and 21 (Fig. 1B & C). Specifically, while *Copg2* expression in ESCs was predominantly biallelic (56% of RNA-seq reads derived from the maternal allele), the maternal allele bias increased to 68.4% by day 7 and reached nearly 73% by day 21 (Fig. 1D). During this period, the two other imprinted genes in the domain, *Klhdc10* and *Klf14*^27^, displayed distinct expression profiles: *Klhdc10* maintained robust expression with a gain of a slight maternal allele bias, whereas *Klf14*, barely detectable in ESCs, exhibited modest maternal induction that peaked at day 7 of differentiation (Supp. Fig. 1C).

Overall, this expression pattern mirrors what is observed in the brains of newborn mice *in vivo*, where *Mest*/*MestXL* is paternally expressed and *Copg2* exhibits a maternal allele bias of 73 % (Fig. 1B, D, and Supp Fig. 1C). These findings underscore the relevance of brain organoids as a model for studying the mechanisms acting at the *Mest* domain, particularly for exploring the induction of *MestXL* and the allelic switch of *Copg2* during brain development.

### *Mest* and *MestXL* Share the Same Promoter Region

The generation of the *MestXL* isoform is thought to be regulated by a brain-specific alternative polyadenylation mechanism^16^. However, given that multiple alternative promoters have been proposed to regulate *Mest* expression^28^, it remains unclear whether this alternative polyadenylation specifically affects the main *Mest* transcript. RNA-seq profiling, conducted at various stages of brain organoid formation and in neonatal brain, revealed no evidence of transcription upstream of the exon overlapping the canonical CpG island/*Mest* promoter, located within the maternally methylated ICR (Fig. 1B). Notably, no transcriptional signal was detected from the upstream promoter previously documented to initiate an oocyte-specific *Mest* transcript^16,28^ (Fig. 1A). Additionally, data mining of long-read RNA-seq performed on adult mouse cerebral cortex^29^ identified isoforms that initiated at the canonical *Mest* promoter and elongated within the *Copg2* genomic region (Supplementary Fig. 2A).

Supporting these observations, Cut & Run experiments in ESCs and neonatal brains, combined with analyses of datasets from ESCs and neural progenitor cells, revealed that only the canonical *Mest* promoter exhibits hallmarks of active transcription (Fig. 2A). In addition to the expected hemimethylation pattern and maternal binding of *Zfp57* in ESCs (characteristic features of an ICR/gDMR region), this active signature includes paternal allele-specific enrichment of H3K4me3 and the presence of ATAC-seq peaks (Fig. 2A). In contrast, the *Mest* upstream region, including the oocyte-specific promoter, is marked by the repressive H3K27me3 modification on its paternal allele (Fig. 2A). During brain organoid formation, ChIP-qPCR analyses demonstrated a progressive increase in the active histone marks H3K9ac and H3K27ac at the canonical promoter starting at day 7, coinciding with peak *Mest* expression and the onset of *MestXL* induction (Fig. 2B). Paternal enrichment for H3K27ac was further observed in the neonatal brain (Fig. 2A). Furthermore, H3K36me3, a histone mark associated with an active transcription elongation, was specifically detected downstream of the canonical promoter, reinforcing the absence of an active upstream promoter for *Mest* in neural tissues (Supplementary Figure 1B).

**Figure 2:**
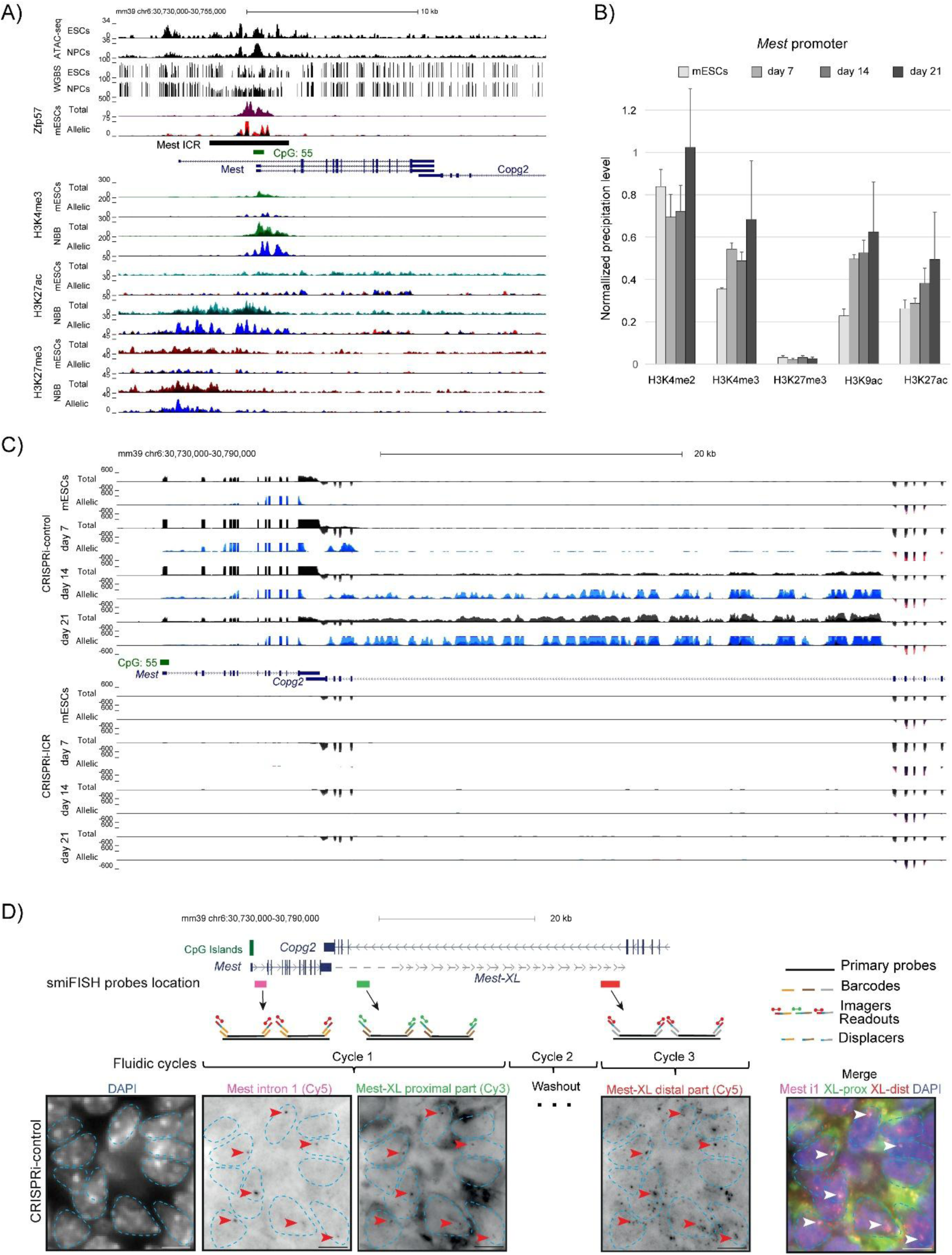
Epigenetic signature and functional analysis of the *Mest* canonical promoter support a shared promoter region with *MestXL*. **A)** Genome Browser view of the *Mest* gene showing CpG island (CGI) positions, ATAC-seq, and methylation (WGBS) data in ESCs and neural progenitor cells (NPCs), as well as ZFP57, H3K4me3, H3K27ac and H3K27me3 enrichment in ESCs and NBB tissues. Data for ZFP57 and histone marks were obtained from B/J material, with quantitative and merged parental allelic signals displayed in the upper and lower panels, respectively. Maternal and paternal enrichments are shown in red and blue, respectively. **B)** Native ChIP-qPCR analysis of histone mark deposition at the *Mest* canonical promoter. Values represent the mean of three independent ChIP experiments (n = 3), each performed in duplicate using B/J ESCs and at days 7, 14, and 21 of in vitro brain organoid differentiation. Precipitation levels were normalized to the *Rpl30* promoter (for H3K27ac, H3K9ac, H3K4me2, and H3K4me3) and the HoxA3 promoter (for H3K27me3). **C)** Genome Browser view of the *Mest* and *MestXL* genomic region showing allelic RNA-seq signals in CRISPRi-control (upper panel) and CRISPRi-ICR (lower panel) B/J ESCs, as well as at days 7, 14, and 21 of in vitro brain organoid differentiation. Values represent the mean of three independent RNA-seq experiments, with quantitative and merged parental allelic RNA-seq signals shown in the top and bottom panels, respectively. Maternal and paternal expression levels are indicated in red and blue, respectively. **D)** Sequential smiFISH experiments with probes targeting the intron 1 of the *Mest* transcript (magenta) and proximal (green) and distal (red) regions of the *MestXL* transcript. The location of probes along the transcript is shown in top panel. Transcripts are targeted with primary probes flanked by two flaps of a unique barcode. Each probe is visualized by sequential hybridization and washout cycles using specific complementary readout and displacer oligos (middle panel). The bottom panel shows representative cells from a day 14 CRISPRi-control brain organoid cryosection. Red arrowheads report the position of the *Mest* nascent transcript (intronic probe) in the different rounds of imaging. Images are maximum intensity projections of 5 planes. Merged images are presented on the right. Nuclei are indicated by blue dashed lines from DAPI images. Scale bars: 5 µm.

To determine whether *Mest* and *MestXL* share the same promoter, we employed two complementary approaches. First, we generated CRISPR inhibition (CRISPRi-ICR) hybrid ESC lines that express dCas9 fused to the transcriptional repressors KRAB and MeCP2^30^ and either a sgRNA control or a sgRNA targeting the canonical promoter of *Mest*, resulting in the transcriptional repression of *Mest*^31^. Second, we generated a hybrid ESC line with a paternal deletion of the promoter region (*Mest* proKO line) (Supplementary Fig 3A). Transcriptional profiling using RNA-seq further demonstrated that repressing the transcriptional activity of *Mest* promoter (using CRISPRi) or deleting *Mest* promoter (*Mest* proKO line) both abolished *Mest* expression in ESCs and prevented *MestXL* induction during brain organoid formation (Fig. 2C; Supplementary Fig. 2 B & 3B). In addition, we performed on brain organoids cryosections sequential single-molecule inexpensive fluorescence in situ hybridization (Seq-smiFISH) experiments^32^, a multiplex technique to visualize and map various RNA species^33^. We designed three sets of probes targeting the *Mest* locus at different positions (Fig. 2D, top). Each set was tagged with a unique barcode^34^. Sequential injections of complementary readout oligos (Fig 2D, central panel) showed that the probes targeting the first intron of *Mest* (1.5 kb, nascent RNA) yielded one single spot per nucleus, confirming its monoallelic expression in brain organoids. Probes targeting proximal (1.6 kb) and distal regions of *MestXL* (2.4 kb) (flanking *Copg2* intron 20), generated signals in proximity to the *Mest* intronic probe, suggesting that the three probes target the same transcript (Figure 2D, bottom panel). Importantly, no signal was observed in the CRISPRi-ICR organoids or after displacer injection and bleaching (Supplementary Fig. 2C & D). Collectively, these data suggest that *MestXL* is an extended form of the *Mest* transcript and establish that both transcripts initiate from the same canonical CpG island promoter defining the *Mest* ICR.

### Maternal switch of *Copg2* is an imprinting event controlled by the ICR

To confirm that the maternal switch of *Copg2* is an imprinting event controlled by the *Mest* ICR, we utilized the CRISPRi-ICR cell line. In this line, repression of the ICR promoter is associated with a gain of DNA methylation on both alleles, effectively “maternalizing” the ICR region (Supplementary Figure 4A). While this cell line retains its differentiation capacity (Supplementary Figure 4B), quantitative RNA-seq analysis revealed that, unlike in wild-type (WT) and CRISPRi-control cells, the predominantly biallelic expression of *Copg2* observed in CRISPRi-ICR embryonic stem cells persisted throughout brain organoid formation (Figure 3A–B). This indicates that the maternal allele bias of *Copg2* does not emerge under these conditions. During this period, overall *Copg2* expression levels remained significantly elevated in CRISPRi-ICR cells compared to CRISPRi-control lines. For instance, by day 21, *Copg2* transcript levels were approximately 30% higher in CRISPRi-ICR cells than in controls (Figure 3C). To further validate these findings, we employed a reciprocal J/B ESC CRISPRi-ICR line for allelic expression analysis. This analysis confirmed that the maternal allele bias of *Copg2* emerged as early as day 7 in the J/B CRISPRi-control line, while consistent biallelic expression was maintained at all stages in the J/B CRISPRi-ICR line. These results provide compelling evidence that this allelic bias is governed by genomic imprinting (Supplementary Figure 4C).

**Figure 3:**
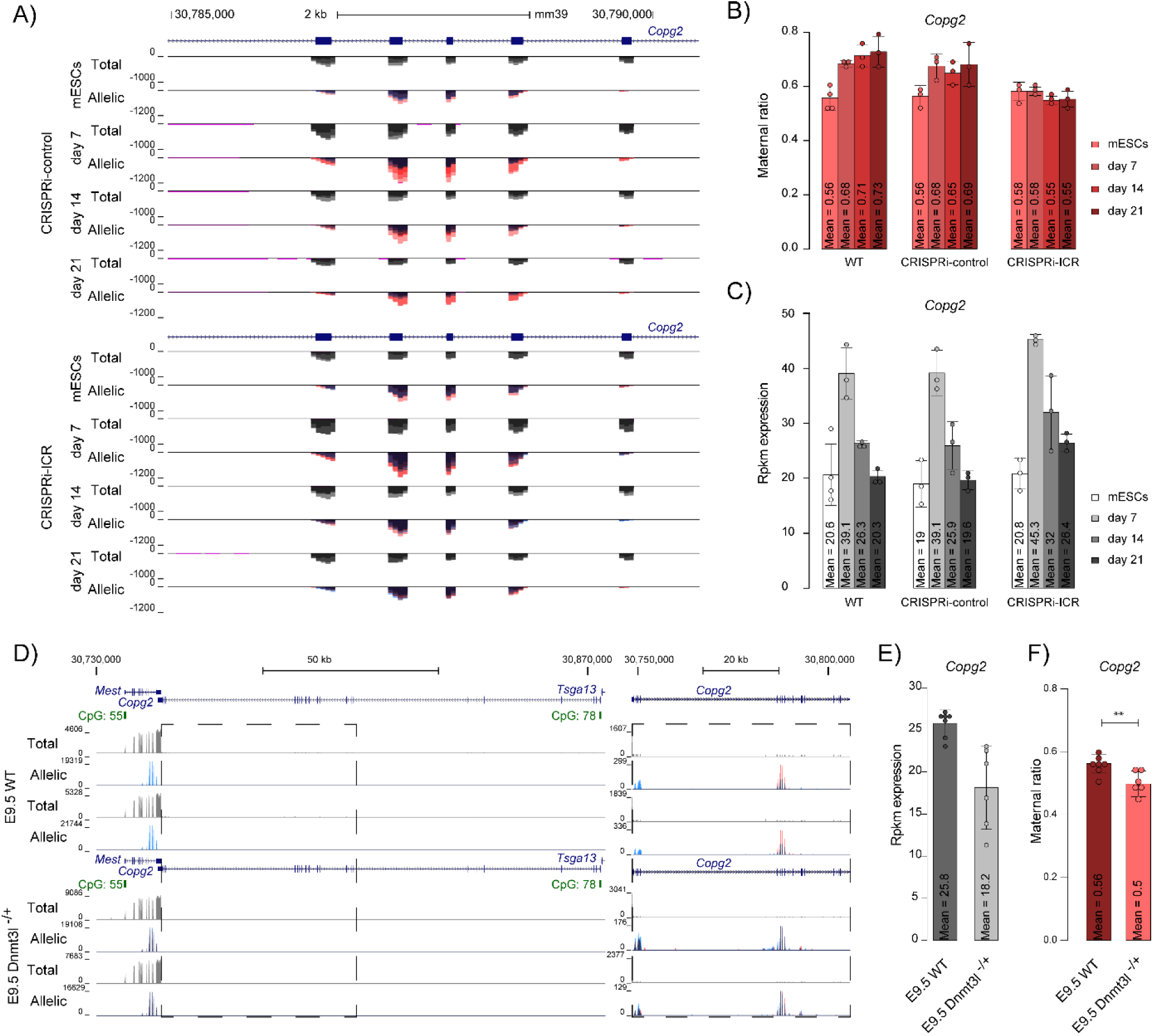
Maternal allele bias of *Copg2* is mediated by the ICR. **A**) Genome Browser view of *Copg2* exon 16-20 showing allelic RNA-seq signals in CRISPRi-control (upper panel) and CRISPRi-ICR (lower panel) B/J ES cells, as well as at days 7, 14, and 21 of *in vitro* brain organoid differentiation. Data represent the overlay of three independent RNA-seq experiments, with quantitative and merged parental allelic signals displayed in the top and bottom panels, respectively. Maternal and paternal expression levels are shown in red and blue, respectively. **B)** Ratio of maternal *Copg2* expression, calculated as the percentage of RNA-seq reads derived from the maternal allele in WT, CRISPRi-control and CRISPRi-ICR ES cells and at days 7, 14, and 21 of brain organoid differentiation. **C)** *Copg2* expression levels, measured as RPKM from RNA-seq, in WT, CRISPRi-ICR and CRISPRi-control ES cells and at days 7, 14, and 21 of brain organoid differentiation. **D)** Genome Browser view of *Copg2* showing allelic RNA-seq signals in representative wild-type (WT) and *Dnmt3L*^-/+^ E9.5 embryos. Quantitative and merged parental allelic signals are displayed in the top and bottom panels, respectively, with maternal and paternal expression levels in red and blue, respectively. **E)** *Copg2* expression levels, measured as RPKM from RNA-seq, in WT (n = 6) and *Dnmt3L*^-/+^ (n = 5) E9.5 embryos. **F)** Ratio of maternal *Copg2* expression in WT (n = 6) and *Dnmt3L*^-/+^ (n = 5) E9.5 embryos, calculated as the percentage of RNA-seq reads derived from the maternal allele. Statistical significance was determined with Mann Whitney test (p < 0.05 *, p < 0.01 **, p < 0.001 ***)

In a complementary approach, we assessed *Copg2* expression in E9.5 Dnmt3l^-/+^ embryos derived from Dnmt3l^-/-^ females, in which maternal DNA methylation imprints at ICRs are not established during oogenesis^35,36^. In wild-type embryos, we confirmed the expected paternal-specific expression of *Mest*, consistent with the known imprinting pattern. Additionally, in line with the prominent contribution of neural tissues at this embryonic stage^37^, we observed a maternal allele bias in *Copg2* expression (Figure 3D, Supplementary Figure 4D).

In Dnmt3l^-/+^ embryos, the “paternalization” of the ICR, resulting from the absence of maternal DNA methylation, led to a marked increase in *Mest* expression, which became biallelic. Conversely, *Copg2* expression was significantly reduced and shifted from a maternal allele bias in wild-type embryos to biallelic expression in Dnmt3l^-/+^ embryos (Figure 3E, F). Notably, *MestXL*, assessed through the expression of the transcript annotation Gm56999, was barely detectable in wild-type embryos but exhibited modest biallelic induction in Dnmt3l^-/+^ embryos (Supplementary Figure 4D).

Together, these observations confirm that ICR-mediated mechanisms are essential for establishing the maternal allele bias of *Copg2*. They further suggest that a maternally methylated ICR promotes *Copg2* expression, whereas an unmethylated ICR tends to decrease it.

### An Allelic Chromatin Architecture at the *Mest* Locus

The prevailing model posits that *MestXL* transcript drives the shift of *Copg2* from biallelic to a maternal-biased expression in neural tissues by interfering with paternal allele transcription in cis^16^. However, allelic expression dynamics observed in brain organoid models reveal a robust maternal allele bias for *Copg2* as early as day 7, a stage at which *MestXL* expression remains minimal (Fig. 1 B &C). This temporal discrepancy raises questions about the nature of the ICR-based mechanism involved in the initiation of the biased *Copg2* allelic usage. Chromatin signature analysis at the *Copg2* promoter showed the absence of the repressive mark H3K27me3 and a progressive gain of activating marks (H3K27ac and H3K9ac) from day 7 onward, aligning with the observed peak of *Copg2* expression (Fig. 4A & B). This chromatin signature pattern at the *Copg2* promoter is consistent with the involvement of an upstream enhancer, potentially acting in an allele-specific manner due to the insulator function of the ICR. If this hypothesis holds, the *Mest* locus would be expected to exhibit a three-dimensional (3D) allelic organisation centred on its ICR. To test this hypothesis, we characterized the higher-order chromatin architecture of the *Mest* locus. We first mapped the genome-wide distribution of CTCF (Fig. 4C). Using allele-specific C&R assays, we confirmed previous findings^11^ that CTCF binds the canonical *Mest* promoter/ICR exclusively on the paternal allele in newborn brain tissue. In contrast, CTCF binding at other sites within the *Mest* domain occurs on both alleles.

**Figure 4:**
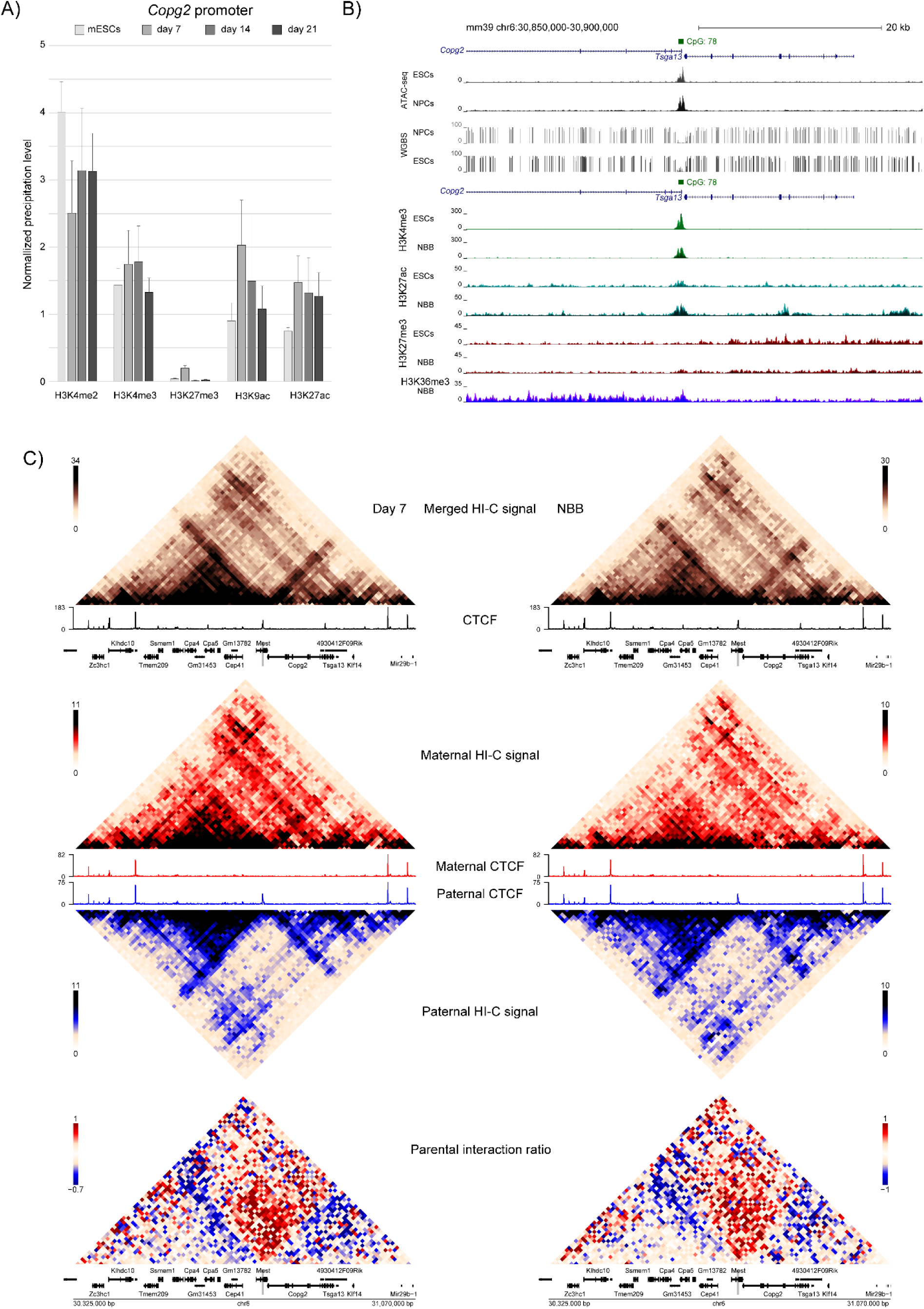
Higher-order chromatin structure at the *Mest* locus. **A)** Native ChIP-qPCR analysis of histone mark deposition at the *Copg2* promoter. Values represent the mean of three independent ChIP experiments (n = 3), each performed in duplicate using B/J ESCs and at days 7, 14, and 21 of *in vitro* brain organoid differentiation. Precipitation levels were normalized to the *Rpl30* promoter (for H3K27ac, H3K9ac, H3K4me2, and H3K4me3) and the *HoxA3* promoter (for H3K27me3). **B)** Genome Browser view of the *Copg2* gene showing CpG island (CGI) positions, ATAC-seq, and methylation (WGBS) data in ESCs and NPCs, as well as H3K4me3, H3K27ac, H3K27me3, and H3K36me3 enrichment in ESCs and NBB tissues. Histone mark data were obtained from B/J material, with quantitative and merged parental allelic signals displayed in the upper and lower panels, respectively. Maternal and paternal enrichments are shown in red and blue, respectively. **C)** Genome Browser view of the *Mest/Copg2* domain showing Hi-C capture-derived contact matrices in B/J day 7 brain organoids (left panel) and NBB (right panel). Merged interactions are shown in brown, maternal in red, and paternal in blue, along with merged (in black) and allelic (maternal: red, paternal: blue) CTCF C&R signals in newborn brain (NBB). The log2 maternal/paternal interaction ratio is displayed in the lower panel.

To assess whether these CTCF-bound regions shape the 3D organization of the *Mest* domain, we performed allele-specific capture Hi-C at day 7 of brain organoid formation and in newborn brain tissue. Using biotinylated probes spanning an 800 Kbp region of the *Mest* imprinted locus and neighboring genes, we found that the *Mest* domain is embedded within a larger chromatin structure, flanked by two biallelic CTCF-binding sites in intergenic regions downstream of *Zc3Hc1* and upstream of *Klf14*. Within this framework, two subdomains, anchored at the canonical *Mest* promoter/ICR, exhibit enhanced chromatin interactions (Fig 4C). These subdomains are present at day 7 and observed in the newborn brain, a pattern we validated by reanalyzing high-resolution, non-allelic Hi-C datasets from in vivo neural progenitor cells (Supp. Fig. 5).

**Figure 5:**
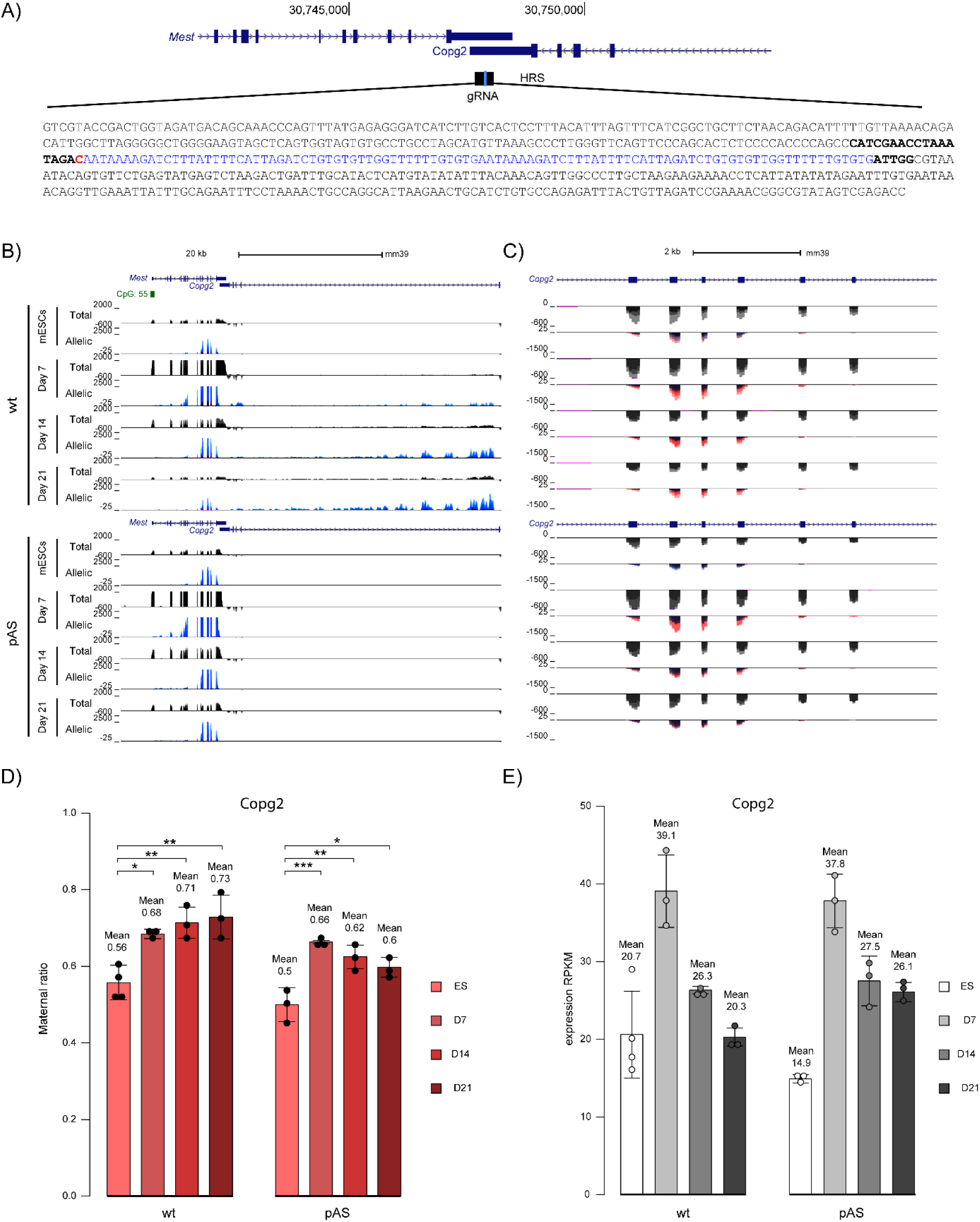
Role of *MestXL* in the maternal switch of *Copg2* during brain organoid development. **A**) Schematic overview of the strategy used to generate the ES-pAS cell line. Double synthetic polyadenylation signals (pAS) are shown in blue (2 × 49 bp). The sequence of the gRNA used is shown in bold, and the cutting site is marked in red. **B)** Genome Browser view of the *Mest/Copg2* locus showing allelic RNA-seq signals in B/J pAS ESCs and B/J WT ESCs and at days 7, 14, and 21 of brain organoid differentiation. Data represent the overlay of three independent RNA-seq experiments, with quantitative and merged parental allelic signals displayed in the top and bottom panels, respectively. Maternal and paternal expression levels are indicated in red and blue, respectively. Zoom on *Copg2* exon 16-20 is shown in **C). D)** Ratio of maternal *Copg2* expression, calculated as the percentage of RNA-seq reads derived from the maternal allele in WT and pAS ESCs (n = 3), as well as at days 7 (n = 3), 14 (n = 3), and 21 (n = 3) of *in vitro* brain organoid differentiation. Statistical significance was determined with the one-way ANOVA and a post-hoc Tukey test (one-way ANOVA p<0.05 and Tukey test p < 0.05 *, p < 0.01 **, p < 0.001 ***). **E)** *Copg2* expression levels, measured as RPKM from RNA-seq, in wild-type (WT) and pAS ESCs (n = 3), as well as at days 7 (n = 3), 14 (n = 3), and 21 (n = 3) of *in vitro* brain organoid differentiation.

To determine the parental origin of these substructures, we leveraged strain-specific SNPs to generate allele-specific contact maps. Striking allelic differences are present at both day 7 and in newborn brain tissue: the two subdomains anchored at the *Mest* promoter/ICR are specifically visible on the paternal allele (Fig 4C). This parental bias in chromatin contacts is further highlighted by maternal/paternal interaction ratio maps (Fig. 4C).

Collectively, these findings demonstrate that allele-specific CTCF binding structures an allelic 3D chromatin organization within the *Mest* domain, providing a structural framework for an allele-specific enhancer insulating function of the *Mest* ICR.

### *MestXL* is dispensable for initiating but essential for maintaining the maternal allelic bias of *Copg2* during brain organoid development

In brain organoids derived from the CRISPRi-ICR cell line, the loss of allelic bias in *Copg2* expression could result from either the absence of *MestXL* or other ICR-mediated mechanisms, such as those related to its insulator function. To specifically assess the role of *MestXL* in the allelic switch of *Copg2*, we used CRISPR-Cas9-mediated recombination to generate a hybrid ES cell line (ES-pAS) with biallelic insertion of two synthetic polyadenylation signals (pAS) in the 3’UTR region of *Mest* (Figure 5A). This insertion preserved the integrity of the *Mest* transcript, which maintained its wild-type expression profile during brain organoid differentiation, including sustained paternal expression (Fig. 5A), a peak expression at day 7 (Suppl Fig. 6A), and the capacity to produce MEST protein, in contrast to the CRISPRi-ICR line (Suppl. Fig. 6B). In contrast, *MestXL* expression was abolished, with no or barely detectable RNA-seq and RT-qPCR signals at any stage (Figure 5B, Supplementary Figure 6A). This further demonstrates that *MestXL* extends from *Mest*. Thus, we next used this novel model to isolate the specific contribution of *MestXL* to the allelic bias of *Copg2*.

In the pAS cell line, as in the WT line, *Copg2* initially exhibited biallelic expression at the ESC stage (Figure 5 C& D). Remarkably, however, by day 7 of brain organoid formation, it acquired a maternal allele bias of 66%, closely matching the 68% bias observed in wild-type (WT) cells, despite the absence of *MestXL*. This suggests that *MestXL* is not required to initiate the maternal allele bias.

Strikingly, while the maternal allele bias in WT cells increased progressively (71% at day 14 and 73% at day 21), it declined in ES-pAS-derived cells, reaching 62% at day 14 and 60% by day 21, (Figure 5C & D). This contrasting trend strongly suggests that *MestXL* plays a dominant, though not exclusive, role in maintening this bias at later developmental stages.This divergence was further reflected in *Copg2* expression levels: at day 7, expression was similar between WT and ES-pAS-derived cells, but by day 21, *Copg2* levels were higher in ES-pAS-derived cells (Figure 5E). This elevated expression may result from the increased stability of the paternal transcript in the absence of *MestXL*.

Collectively, these findings demonstrate that *MestXL*, consistent with its low expression at early stages, is dispensable for initiating the maternal switch of *Copg2* in day 7 neural progenitor-enriched cells. However, it becomes essential for sustaining this bias at later stages, particularly at Day 21. Thus, the role of *MestXL* in regulating the maternal allele bias of *Copg2* is neural stage-specific.

## Discussion

In this study, we investigated the molecular mechanisms underlying the maternal allelic switch of *Copg2* during neural differentiation. By leveraging a stem cell-based brain organoid model, we integrated multi-omic analyses with functional approaches to dissect the regulatory events governing both the induction and maintenance of *Copg2*’s maternal allele bias throughout neural lineage specification. Our findings challenge the established model, which posits that transcriptional interference mediated by the long *Mest* isoform, *MestXL*, is solely responsible for this maternal allele bias. Instead, we demonstrate that the initiation of the allelic switch occurs independently of *MestXL*, revealing that both temporal and neural stage-specific regulatory mechanisms collaborate to finely tune *Copg2* expression during the formation of neural lineages.

The brain organoid model enabled us, for the first time, to examine the developmental timing of *MestXL* induction and the allelic switch of *Copg2* during brain development. Our findings uncover dynamic expression patterns within this genomic domain over the observed time window. Specifically, *Mest* expression peaks at the neural progenitor stage (Day 7), coinciding with the initial induction of *MestXL*, which reaches full production in neuron and neuronal-precursor -enriched (Day 14) and neuron-enriched and beginning of gliogenesis (Day 21) stages. Previous analysis of SAGE tags demonstrated that *MestXL* is a polyadenylated RNA terminating transcription approximately 720 bp upstream of *Copg2* exon 20^16^. Consistent with these data, our results support a model in which *MestXL* is transcribed from the canonical *Mest* promoter. This transcript spans a genomic region of nearly 45 kb, including approximately 38 kb within the *Copg2* gene, encompassing its last four exons and most of intron 20. Notably, a substantial portion of this region is retained in the mature RNA, suggesting a transcript with extensive terminal exons. Strikingly, our data indicate that the annotated gene model *Gm5699*, located at the 3’ end of *MestXL*, previously proposed as a novel paternally expressed gene named *Mit1/Lb9*^38^, as well as several so-called *Copg2* antisense transcripts reported in the literature (*Copg2AS*, *Copg2os2*), all correspond to distinct regions of *MestXL*. This reinterpretation unifies these observations under a single transcriptional unit. Further characterization, for instance through long-read sequencing, will be necessary to fully assess the mature size of *MestXL*. These structural features align with four recently predicted RefSeq *Mest* transcripts (XM_036165899, XM_036165900, XM_036165901, XM_036165902), each spanning nearly 45 kb, whose two terminal exons (approximately 22 kb and 7.5 kb, respectively) cover the last four exons and the majority of intron 20 of *Copg2*, reinforcing the proposed structure of *MestXL* as a large unspliced transcript or with large terminal exons.

We also observed that the maternal switch of *Copg2* is induced as early as the neural progenitor stage, coinciding with an increase in its expression, and is maintained in neuron-enriched stages, where overall *Copg2* expression levels subsequently decrease. This profile demonstrates that the maternal switch of *Copg2* can be decoupled from the presence of *MestXL*. This observation aligns with the central finding of our study: *MestXL* is not required for the induction of *Copg2*’s maternal allelic switch in neural progenitors, although it is necessary for its maintenance at later differentiation stages. This indicates that additional regulatory mechanisms govern the establishment of its maternal allele bias during neurogenesis.

The observed increase in *Copg2* expression at neural progenitor stage, combined with the absence of repressive histone marks and the gain of transcription-activating marks at the *Copg2* promoter, suggests that the appearance of the maternal switch results from enhanced expression of the maternal allele rather than transcriptional repression of the paternal allele. Furthermore, given the peak of paternal *Mest* expression observed in neural progenitors, and the identification of an ICR-anchored allele-specific three-dimensional chromatin organization within this domain, we hypothesize the involvement of an upstream enhancer acting in an parental allele-specific manner, mediated by the insulator function of the ICR on the paternal CTCF-bound allele, to promote paternal *Mest* and maternal *Copg2* expression, respectively.

Together, these data support a working model in which an enhancer and *MestXL* act sequentially during neural lineage formation to finely regulate *Copg2* expression levels. In neural progenitors, the activation of an upstream enhancer, combined with the insulator activity of the ICR, boosts paternal *Mest* and maternal *Copg2* expression, leading to the observed expression peak for both transcripts at Day 7 and to the induction of *Copg2*’s maternal allele bias. Subsequently, in later stages, the overall reduction and/or absence of enhancer activity combined with increased *MestXL* expression, leads to an overall decrease in *Copg2* levels. Despite this reduction, the maternal allele bias is maintained through the destabilization of the paternal *Copg2* transcript (Figure 6).

**Figure 6:**
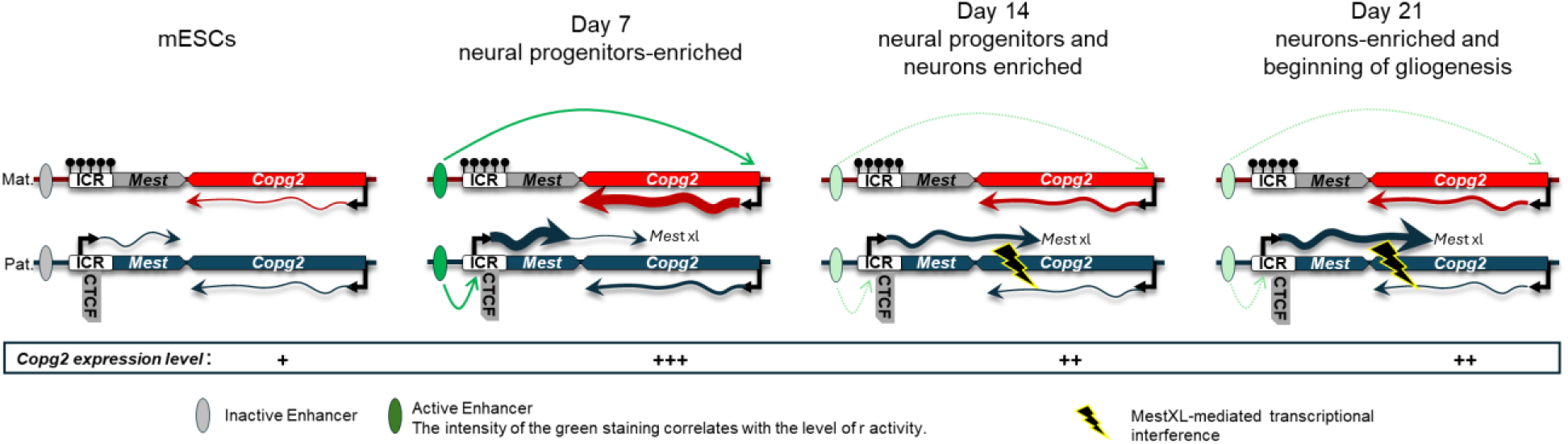
working model of Copg2 regulation during neural differentiation. In ESCs, *Mest* is expressed exclusively from the paternal allele, while *Copg2* is biallelic. As ESCs progress to the neural progenitor stage (organoids at day 7), activation of an upstream enhancer (depicted here as a hypothetical element), together with the insulator activity of the ICR, enhances paternal *Mest* and maternal *Copg2* expression. This results in peak transcript levels for both *Mest* and *Copg2* and the maternal allele bias of *Copg2*. As neurogenesis proceeds (days 14 and 21), the enhancer activity is reduced and/or confined to specific cell types while *Mest* progressively extends into *MestXL*, which induces transcriptional interference (flash sign) that destabilises the paternal *Copg2* transcript, leading to an overall decrease in *Copg2* expression while maintaining its maternal bias. Collectively, the coordinated action of the enhancer, then *MestXL,* finely tunes *Copg2* expression levels and parental bias during neurodevelopment.

It is tempting to speculate that the enhancer-induced gain of *Mest* expression at Day 7 is a prerequisite for *MestXL* formation. RNA polymerase II pausing or reduced processivity downstream of a functional polyadenylation site is known to promote effective polyadenylation^39,40^, whereas enhanced transcription may conversely favour bypass of this site and 3’UTR extension. The initial activation of the enhancer in neural progenitors may therefore exert opposing temporal effects: first, by directly increasing *Copg2* expression, and subsequently, by reducing its levels through *MestXL* formation on the paternal allele at later stages of neural differentiation.

Strong support for this model comes from a recent preprint identifying, through CRISPR interference screening, an enhancer located approximately 150 kb upstream of *Mest* that differentially activates paternal *Mest* and maternal *Copg2* expression during the differentiation of neural precursors into neurons^41^. Further analyses will be required to determine whether this enhancer’s activity is more prominent in neural progenitors, as expected under our model, or whether additional regulatory elements within the domain may also contribute.

Our results reveal a finely tuned and complex regulatory mechanism controlling *Copg2* expression levels, characterized by an upregulation in neural progenitors and a subsequent downregulation at later stages of neurogenesis. *Copg2* encodes the γ2-COP subunit of the COPI complex (Coatomer Protein Complex I), a key player in vesicular transport between the endoplasmic reticulum and the Golgi apparatus. This expression profile suggests an increased demand for *Copg2* and the COPI complex in neural progenitors, where protein synthesis activity is strongly stimulated. Consistent with this, several studies have confirmed that Golgi-ER trafficking, partially regulated by COPI, is essential for maintaining the proliferation, survival, and polarity of neural progenitors, and that disruptions in this trafficking lead to neurogenesis defects and cortical development anomalies^42–45^. In mammals, the γ2-COP subunit has a paralog, γ1-COP (encoded by *Copg1*). Although these paralogs are largely functionally redundant, studies have shown that during retinoic acid-induced differentiation of P19 pluripotent cells into neurons, γ1-COP specifically promotes neurite outgrowth^42^. Our observations, obtained in a brain organoid model, further raise the possibility that γ2-COP may also fulfil a specific role within the COPI complex in neural progenitors. Further studies will be needed to determine whether this transcriptional regulation translates to the protein level, and whether analogous regulation applies to other components of the COPI complex.

Our data indicate that the maternal allele bias observed for *Copg2* is not directly linked to its expression level, but is instead a mechanistic consequence of the regulatory processes at play. This decoupling implies that allelic bias is not functionally determinant per se, but rather reflects broader regulatory mechanisms. From an evolutionary perspective, our results suggest that alternative strategies for fine-tuning *Copg2* regulation, such as distal enhancers bypassing the insulating effect of the ICR, could account for the absence of allelic bias in certain genetic contexts. This provides a coherent framework to explain the contradictory results reported in the literature, where *Copg2* is preferentially expressed from the maternal allele in certain intra-subspecific hybrids but not in interspecific hybrids involving *Mus spretus*^46^.

Our work further demonstrates that *MestXL* is required and represents the primary driver of the maternal allele bias observed for *Copg2* during the late stages of brain organoid formation, particularly at Day 21, a stage enriched in neurons. This observation supports a convergent transcription-based mechanism whereby *MestXL* transcriptionally interferes with the paternal *Copg2* transcript, as seminally proposed by McIsaac et al.^16^. The functional significance of *MestXL* is further underscored by its evolutionary conservation. The *MEST* domain, which includes the convergent orientation of *MEST* and *COPG2*, as well as a maternally methylated DMR in the *MEST* promoter, is conserved in humans and pigs. In these species, *MEST* is paternally expressed, and a paternal antisense transcript (*COPG2IT1*), analogous to mouse *MestXL*, has been identified in brain-derived tissues within intron 20^47,48^. While allelic expression of *Copg2* has not yet been determined in the porcine brain, it is notable that *COPG2* is described as biallelically expressed in the human brain^47^. Given our observations of temporal and neural stage-specific regulation of *Copg2* in mice, further analyses at distinct stages of human neural development, coupled with current approaches capable of assessing allelic expression biases, are required to definitively establish the imprinting status of *COPG2*. Alternatively, the apparent decoupling of *COPG2* and *MESTXL* expression in humans may reflect cell type-specific differences: *COPG2* and *MESTXL* could be expressed in distinct cell types in this species, whereas they co-exist within the same cells in mice. This hypothesis warrants investigation through single-cell transcriptomics and cellular imaging in both mouse and human systems to clarify whether this regulatory mechanism is conserved or has diverged across species.

Taken together, our findings establish that the maternal allele bias of *Copg2* during neural differentiation is not solely driven by *the elongation of Mest transcript into MestXL*, but is instead orchestrated by a two-step regulatory mechanism in which putative enhancer-driven maternal allele activation precedes *MestXL*-dependent paternal allele repression, revealing an unexpected complexity in the regulation of precise dosage of imprinted genes during brain development.

## Material and methods

### Differentiation of mouse ESCs into brain organoids

The hybrid male embryonic stem cell (ESC) lines were previously derived from blastocysts obtained from reciprocal crosses between C57BL/J (B) and JF1 (J) mice^49^ and were maintained in gelatin-coated dishes with ESGRO complete plus medium (Millipore, SF001-500P) containing LIF (leukemia inhibitory factor), BMP4 (bone morphogenetic protein 4), and a GSK3-β (glycogen synthase kinase 3β) inhibitor. For cortical organoid generation, ESCs were dissociated using Accumax and resuspended at a density of 30,000 cells/ml in Cortex Medium I: DMEM/F-12/GlutaMAX supplemented with 10% KSR, 0.1 mM of non-essential amino acids, 1 mM of sodium pyruvate, 0.1 mM of 2-mercaptoethanol, 100 U/mL penicillin/streptomycin, 1 µM DMH1-HCl and 240 nM IWP-2^50^. The media was filtered with a 0.2 µm filter. Cells were seeded into 96-well ultra-low-adhesion plates (Sumitomo) at 3,000 ESCs per well in 100 µL. On day 4, an additional 100 µl of Cortex Medium I was added to each well. On day 7, 4 to 6 aggregates were transferred to 6-well low-adhesion plates and cultured in Cortex Medium 2: DMEM/F-12/GlutaMAX supplemented with N2 and B27 without vitamin A, 500 µg/mL of BSA, 0.1 mM of non-essential amino acids, 1 mM of sodium pyruvate, 0.1 mM of 2-mercaptoethanol, and 50 U/mL penicillin/streptomycin on an agitating platform. On day 14, 2 ml of medium was replaced with 2.5 mL of fresh Cortex Medium 2.

### Material collection

Newborn brains were collected from 1- to 3-day-old mice obtained from reciprocal crosses between C57BL/6J and Mus musculus molossinus JF1. Immediately after dissection, tissues were snap-frozen in liquid nitrogen and stored at −80 °C. The *Dnmt3L* knockout line^36^ was maintained on a mixed 129/terSv x C57BL/6 background. The inbred strain JF1 (Japanese Fancy Mouse 1)^51^ was obtained from the National Institute of Genetics (Mishima, Japan). Embryos at 9.5 days post coitum (dpc) were obtained from the mating of a *Dnmt3L*^+/+^ or *Dnmt3L*^-/-^ female and a JF1 male. Tail DNA was used for genotyping by PCR as previously described^36^.

### DNA methylation analysis

#### DNA extraction and bisulfite sequencing

DNA was extracted as previously described^52^. Bisulfite conversion was performed with the EZ DNA Methylation-Gold Kit from Zymo (ref. D5006), according to the manufacturer’s instructions. PCR amplification, cloning, and sequencing were performed as previously described^52^. Details of the primers used are in Table S2.

#### DNA methylation data mining

ESC and NPC whole-genome bisulfite sequencing (WGBS) data were obtained from GEO DataSets under the accession numbers GEO: <u>GSM748786</u> and GEO: <u>GSM748788</u>, respectively. The reads were first processed using TrimGalore and then mapped to the mm39 mouse genome using Bismark. Duplicate reads were removed with the script deduplicate_bismark. CpG methylation levels were computed from the selected alignments using bismark_methylation_extractor (--no_header --cutoff 4 – bedgraph) and coverage2cytosine scripts. The output was converted with the bedGraphToBigWig tools to be loaded on the University of California, Santa Cruz (UCSC) Genome Browser.

### Expression analysis

#### RNA extraction

RNA was isolated from frozen cell or organoid pellet using TRIzol Reagent (Life Technologies, 15596018) or Zymo Quick RNA Mini Prep (R1055) or Allprep DNA/RNA mini kit (QIAGEN, 80204) according to the manufacturer’s recommendations.

#### RT-qPCR

After treatment with RNase-free DNase I (Life Technologies, 180868-015), first-strand cDNA was generated by reverse transcription with Superscript-IV (Life Technologies, 18090050) or MMLV (Life Technologies, 28025013) using random primers and 500 ng of RNA. Then, cDNA was amplified by real-time PCR with the SYBR Green mixture (Roche) using a LightCycler R 480II (Roche) apparatus. The relative expression level was quantified with the 2-delta Ct method that gives the fold change variation in gene expression normalized to the geometrical mean of the expression of the housekeeping genes *Gapdh*, *Tbp*, and *Gus*. The primer sequences are in Table S2.

For allelic analysis, for each locus of interest, the parental allele origin of expression was assigned following direct sequencing of the cognate RT-PCR product that encompassed a strain-specific SNP (SNP details in Table S2). The relative allelic ratios were quantified from sequence files in ABI format using the Mutation quantification module of the Mutation Surveyor R *_* DNA variant analysis software (Softgenetics, http://www.softgenetics.com/mutationSurveyor.html)

#### RNA sequencing

RNA was collected from WT cell lines (triplicates from the B6×JF1 genetic backgrounds), pAS cell lines (triplicates from the B6×JF1 background), and CRiSPi-ICR cell lines (triplicates from the B6×JF1 background), both at the ES stage and during differentiation into brain organoids at days 7 (D7), 14 (D14), and 21 (D21). RNA was also collected from newborn brains samples lines (from the B6×JF1 and JF1xB6 genetic backgrounds).These samples were used to generate paired-end RNA sequencing (RNA-seq) data. RNA-seq libraries were prepared with the NEBNext Ultra II mRNA-Seq Kit or the Watchmaker mRNA Library Prep Kit and sequenced on an HiSeq4000, NovaSeq6000 or NovaSeq X Plus apparatus by IntegraGen according to the manufacturer’s protocol. To determine the global and allelic expression, RNA-seq reads were mapped to the mm39 genome, masked for JF1 SNPs, using Hisat2 and the list of known splice sites produced with the GENCODE vM37 basic annotation (version 2.2.1; -no-softclip -known-splicesite-infile) . Alignments were filtered with samtools for mapping quality, reads mapped in proper pairs and duplications (version 1.21; view options: -f 2 -q 20, markdup -r). The global alignemnts obtained were treated using SNPsplit to separate allele-specific alignments (version 0.4.0., --paired --no_sort --snp_file JF1 SNPs). The strand-specific coverages of the RNA-seq data were generated using the C57BL/6 and JF1 specific alignments and the global alignments with bamCoverage (version 3.5.6; --normalizeUsing RPKM –filterRNAstrand forward/reverse) and visualized on the UCSC genome browser. Replicates were overlaid for allelic and strand-specific coverage using the track collection builder tool for genome exploration. The read counts per gene were computed for the C57BL/6 and JF1 specific alignments and the global alignments using htseq-count with the GENCODE vM37 basic annotation (version 2.0.9, options: -m intersection-nonempty --additional-attr=gene_name -s reverse). Maternal ratios were computed, only for genes with at least 10 allelic reads, by dividing the number of maternal reads by the sum of allelic reads: (C57BL/6 counts /(C57BL/6 counts + JF1 counts)).

Sequencing libraries for RNA collected from *Dnmt3l*^−/+^ and *Dnmt3l*^+/+^ mouse embryos were prepared using SMART-Seq mRNA LP Kit (TaKaRa, 634768) and Unique Dual Index Kit (Takara, 634756) according to the manufacturer’s protocols. Briefly, 100 ng of total RNA was subjected to cDNA synthesis and amplification by six cycles of PCR. The resultant double-stranded cDNA was enzymatically fragmented and ligated with stem-loop adapters. The adapter-ligated DNA was amplified by 13 cycles of PCR using primers with unique dual indexes to generate Illumina-compatible libraries. The libraries were sequenced using MGIEasy Universal Library Conversion Kit (App A) (MGI, 1000004155) and DNBSEQ-T7RS High Throughput Sequencing Reagent Kit V3.0 (MGI, 940-000268-00) on a DNBSEQ-T7 platform (MGI) with the paired-end mode of 150 bp x2. Due to the complex genetic background of the *Dnmt3l*^−/+^ and *Dnmt3l*^+/+^ embryos, ((129/terSvxC57BL/6)/(JF1)) only SNPs specific to JF1 strain could be used to determine parental origin in RNA-seq experiments. SNPs for the 129Sv and JF1 strains were retrieved from a multisamples VCF produced on the mm39 genome (https://ftp.ebi.ac.uk/pub/databases/eva/PRJEB53906/mgp_REL2021_snps.vcf.gz) and a R script was used to exclude from the JF1 SNPs list those that are common to the 129Sv strain. Next, these filtered JF1 SNPs and the mm39 genome were processed using the bedtools maskfasta tool (version 2.31.1) to generate a masked genome for the SNPs specific to the JF1 background. Paired-end reads were mapped to this masked mm39 genome using Hisat2 and the list of known splice sites produced with the GENCODE vM37 basic annotation (version 2.2.1; options: -no-softclip -known-splicesite-infile). The RNA-seq alignments were filtered with samtools for mapping quality, reads mapped in proper pairs and duplications (version 1.21; view options: -f 2 -q 20, markdup -r). The global alignemnts were treated using SNPsplit in order to separate allele-specific alignments (version 0.4.0., options: --paired --no_sort --snp_file JF1 SNPs). Global and allelic coverage of the RNA-seq were generated using bamCoverage (version 3.5.6; options: -normalizeUsing RPKM) and global and allelic read counts per gene were computed using htseq-count (version 2.0.9, options: -m intersection-nonempty --additional-attr=gene_name -s no). Maternal ratios were computed as previously described.

Total RNA collected from *Mest* proKO cell line was used to generate RNA-seq libraries using “Illumina Stranded mRNA Prep Kit” and sequenced on NextSeq 2000. RNA-seq reads were mapped to the mm39 genome and allelic tracks were created using MEA^53^.

#### LongReads data mining

long-read sequencing from mouse cortex were obtained from SRA database under the project number: PRJNA663877. The Iso-seq reads produced on 8 months cortices (SRR12653624, SRR12653626, SRR12653628) were aligned to the mm39 genome using minap2 (version 2.27-r1193, options -ax splice:hq). Alignemnts were merged and then processed using IsoQuant to identify new transcript isoforms (version 3.4.1, options --stranded none --polya_requirement always --report_novel_unspliced false --data_type pacbio).

### Chromosome conformation analysis

#### Hi-C capture

Hi-C libraries from newborn brains (duplicates from the B6×JF1 background) and brain organoids at days 7 (duplicates from the B6×JF1 background) were generated as previously described in Hi-C 3.0 protocol^54^ with some minor modifications. 5.10^6^ frozen nuclei were used as starting material and were fixed with 1% formaldehyde solution for 10 min. Chromatin digestion was done with 600U of DpnII enzyme during 20 hours at 37°C. After final library PCR amplification, around 1.5 µg of multiplexed Hi-C libraries were mixed equimolarly and concentrated to meet the requirements of the capture. Capture was done with the SureSelect XT HS2/HS technology from Agilent. Agilent SureSelect custom probes targeting Mest locus (chr6: 30300000-31100000 (mm39)) were designed at the ends of DPNII fragments and probes overlapping B6/JF1 SNPs were designed to perfectly match both genotypes. Capture and final amplification were performed with SureSelect XT HS2 Target Enrichment Kit ILM Hyb Module (#5191) according to the manufacturer’s recommendations. Sequencing of Hi-C libraries was performed on an Illumina NovaSeq X Plus apparatus by IntegraGen. Hi-C reads were aligned on the mm39 genome, masked for JF1 SNPs, using HICUP (version 0.9.2). Hi-C alignments were treated using SNPsplit in order to separate parental-specific Hi-C alignments (--hic --snp_file JF1 SNPs). Hi-C alignments assigned to the B6 background for one or both reads were combined using samtools to build the maternal Hi-C alignments. The ones assigned to the JF1 background were used to build the paternal Hi-C alignments. The maternal and paternal specific Hi-C alignments and the global Hi-C alignments were treated using hicup2juicer from HICUP and Pre from Juicer tools to save Hi-C contact matrices in.hic format. Data from replicates were merged to produce the final allelic and global contact matrices. Parental interaction ratio was computed by doing the log2 of the ratio of maternal to paternal signal for the contact matrix at 10 kb resolution with VC_SQRT normalization. Visualization of contact matrices at 10 kb resolution with VC_SQRT normalization was conducted in R (4.5.2) using plotgardener (1.16.0).

#### Hi-C data mining

Hi-C data for NPC purified in-vivo from E14.5 neocortex were obtained from the GEO: GSE96107 DataSet. The 4 replicates were combined and aligned to the mm39 genome using the HiC-Pro (version 3.1.0). The allValidPairs produced by HiC-Pro were treated with the script hicpro2juicebox to save Hi-C contact matrix in .hic format. The contact matrix was visualized at 10 kb resolution with VC_SQRT normalization using plotgardener.

### Cell Imaging

#### Fixation, cryoprotection and sectioning of brain organoids

Organoids were fixed in 4% fresh PFA for 15 min at RT and washed three times for 10 min in PBS. Samples were cryoprotected by incubation in 30% sucrose overnight at 4°C. Organoids were then incubated for 15 min at 37°C and embedded in 7.5% gelatine and 30% sucrose. Blocks were frozen on dry ice and stored at −80°C before sectioning. 10 µm cryosections were done at -18°C using the Cryostat CryoStar NX70DV and directly placed on Superfrost slides (Epredia J1800AMNZ) for immunofluorescence or coated coverslips (see below) for Seq-smiFISH experiment.

#### Immunofluorescence on brain organoid cryosections

Cryosections were air-dried for 5 minutes, rinsed in PBS, blocked and permeabilised in PBS with 0.25% Triton X-100 and 4% horse serum for one hour at RT. Primary antibodies (see Table S) were incubated overnight at 4°C. Cryosections were washed three times with PBS and incubated with secondary antibodies (anti-mouse or anti-rabbit coupled to Cy2 or Cy3 with minimal cross-species reactivity, Jackson Immunoresearch) for 2h at RT, both diluted in PBS containing 0.1% Triton X-100 and 4% FBS. Nuclei were stained with DAPI. The slides were mounted with homemade mowiol and observed under a fluorescent microscope with an Apotome mode (AxioImager Z1, Zeiss) or a confocal microscope (LSM980 AIRYSCAN, Zeiss).

#### Seq-smiFISH

Seq-smiFISH probes against the 3 regions of interest of the Mest locus are listed in Table S2. Primary probes were designed using the oligostan routine^32^ and appended both at 5’ and 3’ end with a unique barcode for sequential imaging detection^34^ Specifically, we used barcode 10 for Mest intron 1, 11 for *Mest-XL* proximal, and 34 for *Mest-XL* distal. FISH probes were purchased from Integrated DNA Technologies at a scale of 50 pmol/oligo as individual oligo-pools and dissolved in 50 µl of TE buffer (10 mM Tris HCl pH 8.0, 1 mM EDTA). Auxiliary oligos (readouts, displacers, and imagers) were purchased from Integrated DNA Technologies at 250 nmole scale and used at 100 µM concentration.

On day 1, 40 mm round coverslips were first washed in isopropanol for 10 min in an ultrasound bath, extensively rinsed with MilliQ water, air dried using compressed air, then coated with 300 µL of Cell-Tak solution (285 µL of 7.5% Na Bicarbonate, 10 µL of Cell-Tak (Corning CB-354240), 5 µL of 1N NaOH), modified from^55^. Coverslips were finally rinsed with DNase/RNase free water (Invitrogen 10977) and air dried for 5-10 min. Cryosections were directly placed on the treated coverslips, fixed in 4% fresh PFA (Electron Microscopy Science 15714S) for 15 min at RT, then rinsed with PBS and permeabilized in 70% ethanol for 1 h at 4°C.

Cryosections were rinsed once and then washed twice for 10 min at RT with Wash I buffer (2X SSC, 1:10 dilution in DNase/RNase free water from filtered 20X SSC, Invitrogen AM9763). Next, they were rinsed once and equilibrated at RT for at least 10 min with Hybridization buffer (2X SSC, 10% formamide (Sigma 47671) in DNase/RNase free water).

Finally, cryosections were incubated overnight at 37°C placed upside-down in a parafilm-coated humid chamber. Hybridization mix consisted of 140 µL of 2X SSC, 10% formamide, 10% dextran (MW > 500000, Sigma 8906), 0.2X NEBuffer3 (New England BioLabs B7003S), 1 mg/mL tRNA (Sigma R8759), 10% fresh Triton X-100 (Sigma T8787), and 0.25 µL of each primary probe.

On day 2, cryosections were rinsed once and washed twice with Hybridization buffer for 30 min at 37°C. Next, they were stained with DAPI (1:1000) for 15 min at RT and placed in Wash I buffer until they were mounted in a FCS2 chamber (Bioptechs 060319-2) for sequential imaging. 40 µL of Readout mix (1X NEBuffer3, 1 µL/readout, 1.05 µL/imager) were heated at 65°C, 1500 rpm for 10 min and let slowly cool down to RT. The first mix was prepared using Mest intron 1 and *Mest-XL* proximal readouts oligos in combination, respectively, with Cy5 and Cy3 imagers. A second mix was prepared to detect *Mest-XL* distal probes using the Cy5 imager. Readout mixes were then diluted in 1.81 mL of Hybridization buffer supplemented with 0.1% Triton X-100 and 1:1000 DAPI. The same buffer was used to prepare the corresponding displacer oligos mixes using 2 µL/displacer in 1.85 mL of buffer. The 4 mixes were then loaded in a 96-well plate. Sequential imaging was performed using a Vutara VXL (Bruker) imaging station integrated with a liquid handling unit (Bruker PlexFlo96). Fluidics cycles alternated readout and displacer mixes and were followed by a round of imaging. Fluidics cycles consisted of sample equilibration with Hybridization buffer (2 mL at 200 µL/min), readout or displacer mix injection (1.6 mL at 200 µL/ min) and incubation, 30 min at 37°C. Next Imaging buffer (2X SSC, 50 mM Tris HCl pH 8.0, 10% D-(+)-Glucose (Sigma G7021), 40 µg/mL catalase (Sigma C40), 0.5 mg/mL Glucose oxydase (Sigma G2133) was injected (2 mL at 200 µL/min) and let equilibrate for 2 min. In the case of displacer cycles, an extra step of sample photobleaching (1 min, at maximum power laser) was performed before flushing the imaging buffer. Images were then acquired at different selected positions (15 to 25 locations/organoid, 2 to 3 organoids/experiment) with 250 nm z-stack and 100 ms exposure time.

#### Western Blotting

Seven D7 organoids were pooled, rinsed with PBS 1x, and kept at -80°C until processing. Pooled organoids were resuspended in 120 µl Laemmli 1X (containing 1.4M beta-mercaptoethanol), disrupted through passing through a Hamilton syringe, and heated for 5 min at 65°C. 20 µl of cell lysates were resolved by SDS-PAGE, blotted onto nitrocellulose membrane and probed overnight at 4°C with an antibody to MEST (Rabbit mAb, ABCAM, ab151564). The nitrocellulose membrane was then incubated with an anti-rabbit antibody conjugated to horseradish peroxidase (HRP) for 60 minutes and developed using the Trident Femto Western chemiluminescent HRP substrate (Genetex) on a Chemidoc apparatus (Biorad). The membrane was reblotted with ACTIN (mouse, Chemicon, MAB1501) to check for equal loading.

### Chromatin immunoprecipitation

#### ChIP-qPCR

Chromatin immunoprecipitation (ChIP) of native chromatin was performed as described by Brind’Amour et al.^56^ using 500,000 cells per immunoprecipitation. Results presented in this article were obtained from at least three ChIP assays performed using independent chromatin preparations, as indicated in the figure legends. Details of the antisera used can be found in Table S3. Quantitative and allelic analyses were performed as described previously in Maupetit-Méhouas et al.^57^. Details of the SNPs and primers used can be found in Table S2.

#### Cut&Run

Cut&Run (C&R) was performed using the CUTANA CUT&RUN Kit (Epicypher) and non-fixed nuclei from ESCs (duplicates from the B6×JF1 background) and newborn brain samples (duplicates from the B6×JF1 and JF1xB6 background), according to the manufacturer’s instructions. The antisera used are listed in Table S3. Briefly, nuclei were isolated from fresh ESCs or frozen newborn brain samplesand stored in nuclear extraction buffer at −80°C. After thawing, 500,000 nuclei per reaction were aliquoted and incubated with pre-activated concanavalin A-coated beads at room temperature for 10 min, followed by overnight incubation with 0.5 μg of antibody in buffer containing 0.01% digitonin at 4°C. Then, nuclei bound to concanavalin A-coated beads were permeabilized with a buffer containing 0.01% digitonin and incubated with the pAG-MNase fusion protein at room temperature for 10 min. After washing, chromatin-bound pAG-MNase cleavage was induced by the addition of calcium chloride to a final concentration of 2 mM. After incubation at 4°C for 2 h, the reaction was stopped by the addition of stop buffer (containing fragmented genomic *Escherichia coli* DNA as spike-in). Following fragmented DNA purification, Illumina sequencing libraries were prepared from ∼5 ng of purified DNA using the CUTANA CUT&RUN Library Prep Kit (EpiCypher 14–1001 and 14–1002) according to the manufacturer’s recommendations. Purified multiplex libraries were diluted to 9 nM concentration (calculated with the Qubit dsDNA HS Assay Kit) and sequenced on a NovaSeq X Plus instrument (Illumina) by IntegraGen SA. Paired-end reads were mapped to the mm39 masked genome and hybrid genome using Bowtie2. Alignment filtering was done with samtools (view -f 2 -q 20), and the coverage was obtained with bamCoverage (global coverage --normalizeUsing RPKM --binSize 25; allelic coverage: --binSize 25). The UCSC track collection builder tool was used to overlay allelic coverages for genome exploration..

#### ATAC-seq data mining

Assay for transposase-accessible chromatin using sequencing (ATAC-seq) data for ESCs and ESC-derived NPCs were obtained from the GEO: <u>GSE155215</u> DataSet. Paired-end reads were treated with trim_galore (--paired) and then were aligned to the mm39 genome using bowtie2 (--very-sensitive -X 1000). Only properly paired alignments were conserved with samtools (view -f 2) and alignments to mitochondrial sequences and random chromosomes were excluded. PCR duplicates were removed using picard-tools (MarkDuplicates --REMOVE_DUPLICATES true). The coverage was assessed using bamCoverage (--normalizeUsing RPKM --binSize 20) and was visualized on the UCSC genome browser. Peaks were called using macs2 (callpeak -f BAMPE --broad --broad-cutoff 0.05 --keep-dup all).

#### CRISPR-Cas9 inhibition

Hybrid B/J and J/B ESC CRISPRi expressing dCas9-KRAB-MeCP2 and either a control sgRNA or a sgRNA targeting the Mest-DMR (ICR) were generated as previously described for E14 ESC CRISPRi control and Mest lines^31^.

#### CRISPR-Cas9-mediated pAS insertion

The guide RNA (gRNA), designed using the ChopChop web tool (https://chopchop.cbu.uib.no/), and the 559-bp homology-directed repair (HDR) template (see supplementary Figure 6), containing two tandem copies of a synthetic 49-bp polyadenylation signal (5′-AATAAAAGATCTTTATTTTCATTAGATCTGTGTGTTGGTTTTTTGTGTG-3′) derived from the rabbit β-globin pAS^58^, were synthesized by Integrated DNA Technologies (IDT). For pAS insertion, 1 × 10⁶ B6/JF1 embryonic stem (ES) cells were nucleofected with 60 pmol of Cas9 protein, 72 pmol of gRNA, and 500 ng of the HDR template, in the presence of an apoptosis inhibitor (Thermofischer Revitacell Supplement #A2644501) and 1 µM HDR enhancer (IDT), using the Mouse Embryonic Stem Cell Nucleofector Kit (Lonza VPH-1001) with program A24. Following FACS cells sorting and single-cell plating into 96-well plates, targeted integration was assessed by colony PCR. Clones harboring the pAS insertion were identified, and one clone displaying biallelic insertion at the target locus (designated ES-pAS) was selected for further characterization.

#### Generation and characterization of F1 hybrid ESCs with a Mest promoter deletion (Mest proKO allele)

The deletion of the Mest promoter region was engineered in the male F1 hybrid ESC line BC1.3 (C57BL/6J female x CAST male) by CRISPR-Cas9 genome editing. The establishment of those ESCs from a F1 hybrid blastocyst and the details of the CRISPR-Cas9 experiment will be published elsewhere (Ha et al., manuscript in preparation). For the deletion, we used two guide RNAs targeting the Mest promoter region: sgRNA5-1 (5’-GCCCACCCCATGGCGGGATC-3’) and sgRNA3-1 (5’-CTAGATCGTGCCGCGCAGTG-3’), each followed by the PAM 5’-TGG-3’ on the (+) strand. After transfection of BC1.3 cells, positive clones were first screened using a genomic PCR reaction spanning the targeted region (5F1 5’- CCTTCCTCCCTCCCCTTAAT-3’ and 3R4 5’-AGTGCCAGAACACGATGGTT-3’). For the heterozygous ESC clone BC11, the structure of the deletion allele was analysed by Sanger sequencing of the 864-bp 5F1-3R4 PCR product with 3F2 (5-ACTAAAGCTGAGAATGGCTGG-3’). This confirmed the deletion of a 2131-bp sequence (GRCm39/mm39 - chr6:30,736,675-30,738,805) of the Mest promoter region on the paternal CAST allele, based on SNP calls at three heterozygous BC SNPs (rs39519970, rs37511830, and rs36857888). A similar analysis of the WT allele using primers internal to the deletion confirmed its C57BL/6J origin. The genotype of ESC line BC11 is therefore Mest+/proKO. The deleted region encompasses the upstream CTCF binding site, and the entire promoter CGI, which includes exon 1 of Mest.

#### Statistical analyses

Statistical analyses were performed using GraphPad or R. The statistical test used for each comparison and the number of independent experiments are indicated in the figure legends.

## Acknowledgments

We thank all members of P.A. and Journot lab for helpful discussion and critical reading of the manuscript. We thank members of the iGReD Bioinformatics Platform (BIM) for their technical assistance and discussions on NGS traitements, Edouard Bertrand (IGH) for helpful discussions and for sharing resources and Julien Béthune for helpful discussion on the coatamers’ function. We acknowledge the imaging facility MRI, a member of the national infrastructure France-BioImaging (https://ror.org/01y7vt929) supported by the French National Research Agency (ANR-24-INBS-0005 FBI BIOGEN) for access to microscopes. We thank Szi Kay Leung and Jonathan Mill for helping us access their long-read RNA-seq data. This research has been financed by the French government IDEX-ISITE initiative 16-IDEX-0001 (CAP 20–25) (Projet Emergence [to P.A. and F.C.] and project Challenge 3-recherche [P.A.]), Fondation pour la Recherche Médicale (FRM MND 2023 to F.C) and Agence Nationale de la Recherche (ANR-23-CE12 CORGI to T.B. and P.A.). B.M is supported by a postdoctoral fellowship (I-SITE Clermont Auvergne Project - European Centre for Health and Human Mobility). Work in the Lefebvre laboratory was supported by a Project Grant (#PJT-165992) from the Canadian Institutes of Health Research (CIHR) and by a Natural Sciences and Engineering Research Council of Canada (NSERC) Discovery Grant (RGPIN-2024-04175).

## Author contributions

Conceptualization: T.B., F.C. and P.A.; Experimental design: S.P., A-C. F., L.L., T.B., F.C. and P.A; Data production: S.P., A-C.F, C.G.G., O.C, M.E.; A.M.; S.P., L.F., C.V-B., D.N., A.H., N.A., H.K., N.G., L.L., T.B and F.C; Data analysis, S.P., A-C.F, C.G.G., K.H, B.M., K.N., L.L., A.B.B., T.B;, F.C and P.A.; Writing – original draft, P.A.; Figure original draft: FC Writing/Figure – review & editing, B.M., L.L., T.B., F.C. and P.A. Supervision, T.B., F.C. and P.A; Project administration, T.B. and P.A.; Funding acquisition: L.L., T.B., F.C. and P.A. All authors read and approved the final manuscript.

---

**Supplementary Figure 1:**
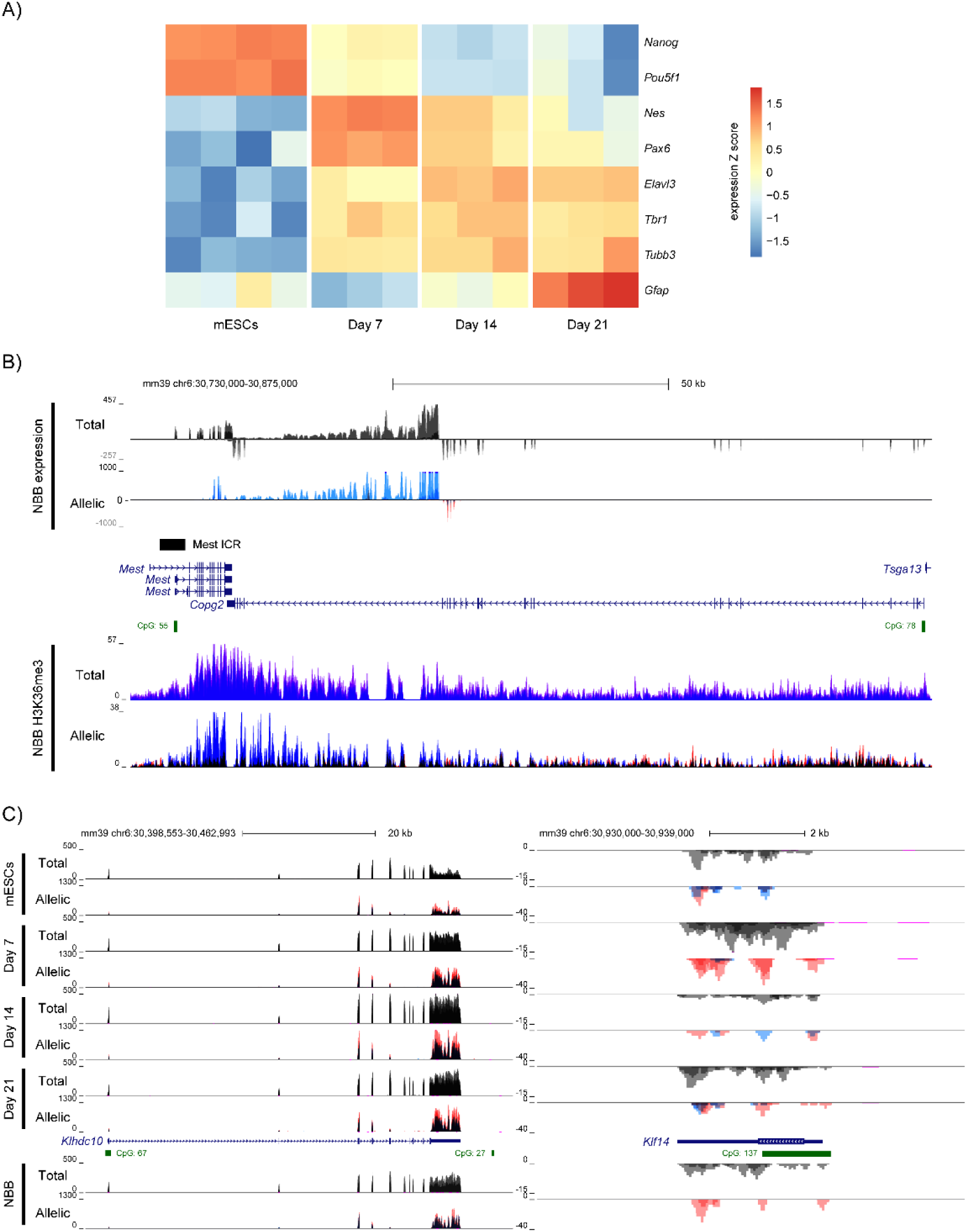
Differentiation and *Mest* imprinted domain dynamics during mouse brain organoid development. **A)** Heatmap illustrating the expression levels of specific markers, as measured by RT-qPCR, during the differentiation of brain organoids. Pluripotency markers (*Nanog*, *Pou5f1*), neural progenitor markers (*Nestin*, *Pax6*), neuronal markers (*Elavl3*, *Tbr1*, *Tubb3*), and the astrocyte marker *Gfap* were analyzed across B/J embryonic stem cells (ESCs) and at days 7, 14, and 21 of brain organoid differentiation. Data are presented for each independent experiment, with sample sizes of n=4 for ESCs and n=3 for days 7, 14, and 21. The color scale represents Z-score expression levels. **B)** Genome browser view of the *Mest* and *Copg2* loci, showing allelic RNA-seq signals (upper panel) and H3K36me3 enrichment obtained by Cut&Run (lower panel) in merged B/J and J/B newborn brain tissue (NBB)(*n* = 2 for RNA-seq; *n* = 4 for C&R). For each condition, the quantitative and merged parental allelic signals are at the top and bottom, respectively. Maternal and paternal expression levels are shown in red and blue, respectively. **C)** Genome browser view at the *Klhdc10* and *Kfl14* genes, respectively, to show the allelic-oriented RNA-seq signals in B/J ESCs (n=4) and at day 7 (*n* = 3), day 14 (*n* = 3), and day 21 (*n* = 3) of in vitro brain organoid differentiation, as well as in merged B/J and J/B newborn brain tissue (NBB) (*n* = 2). For each condition, the quantitative and merged parental allelic RNA-seq signals are at the top and bottom, respectively. Maternal and paternal expression levels are shown in red and blue, respectively.

**Supplemental Figure 2:**
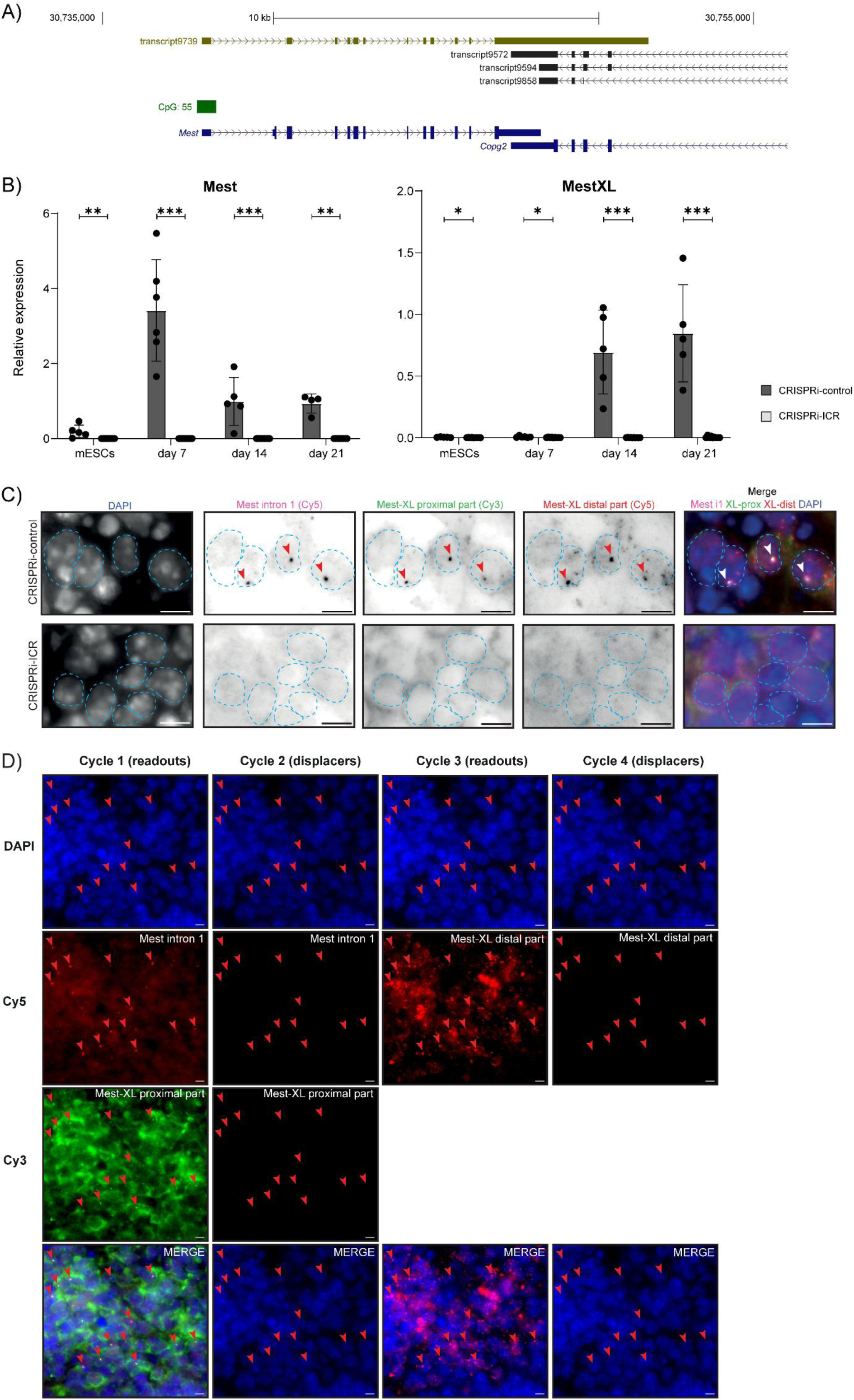
Alteration of the *Mest* canonical promoter eliminates *MestXL* expression in brain organoids. **A)** Genome Browser view of the *Mest* and *Copg2* genes displaying long-read RNA-seq signals in adult mouse cortex. **B)** RT-qPCR analysis of *Mest* and *MestXL* expression levels in CRISPRi-ICR and CRISPRi-control B/J ESCs, as well as at days 7, 14, and 21 of *in vitro* brain organoid differentiation. Expression is presented as a percentage relative to the geometric mean of three housekeeping genes (*Gapdh*, *Gus*, and *Tbp*), with data shown as mean ± SEM. Statistical significance was determined with Mann Whitney test (p < 0.05 *, p < 0.01 **, p < 0.001 ***). **C)** Sequential smiFISH experiments in CRISPRi-control (top) and CRISPRi-ICR (bottom) day 14 brain organoid cryosection with probes targeting the intron 1 of the *Mest* transcript (magenta) and proximal (green) and distal (red) regions of the *Mest-XL* transcript. Red arrowheads report the position of the Mest nascent transcript (intronic probe) in the different rounds of imaging. Images are maximum intensity projection of 5 planes. Merged images are shown on the right. Nuclei are indicated by blue dashed lines from DAPI images. Scale bars: 5 µm. **D)** Washout cycles of sequential smiFISH experiments in CRISPRi-control day 14 brain organoid cryosection. Cycle 1 shows the visualization of *Mest* intron 1 (red, Cy5) and *MestXL* proximal (green, Cy3) readouts that are displaced and bleached in cycle 2 (note the absence of signal). Cycle 3 shows the visualization of *MestXL* distal (red, Cy5) readout that is displaced and bleached in cycle 4. Red arrowheads report the position of the Mest nascent transcript (intronic probe) in the different rounds of imaging. Images (maximum intensity projection of 5 planes) are shown using the same intensity scale. Scale bars: 5 µm.

**Supplemental Figure 3:**
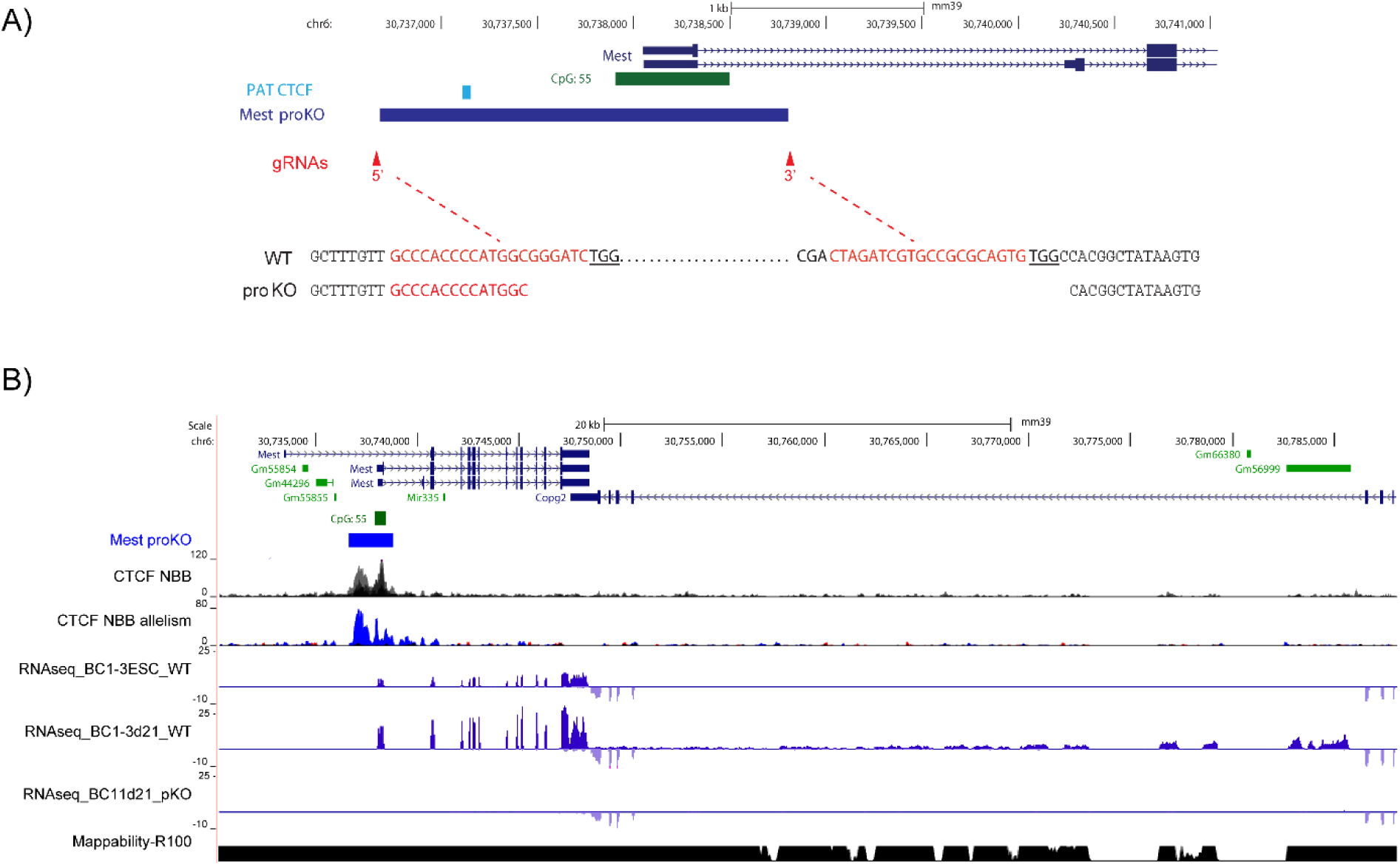
Deletion of the *Mest* canonical promoter eliminates *MestXL* expression in brain organoids. A) Top- Screenshot of a UCSC Genome Browser window from mouse genome build mm39 showing the 5’ end of the *Mest* gene with its exon 1-associated CpG island, the position of the upstream CTCF binding site on the paternal allele and the extent of the 2131-bp *Mest* promoter deletion (*Mest* ProKo), defined by unique 5’ and 3’ gRNA for CRISPR-Cas9 deletion. Bottom-Alignment of the DNA sequences of the WT and *Mest* proKO line alleles, with the position of CRISPR gRNA sequences shown in red and PAMs in bold/underlined characters. B) UCSC Genome Browser session centered on the *Mest-Copg2* region, annotated with BED tracks as in A, and showing: i) CTCF C&R tracks from newborn brains (NBB) with quantitative and merged parental allelic signals displayed in the upper and lower panels, respectively. Maternal and paternal enrichments are shown in red and blue, respectively.; ii) bulk RNA-seq tracks for ESCs and d21 organoids from the WT parental line BC1.3, and d21 organoids from the Mest+/proKO ESC BC11 (pKO).

**Supplemental Figure 4:**
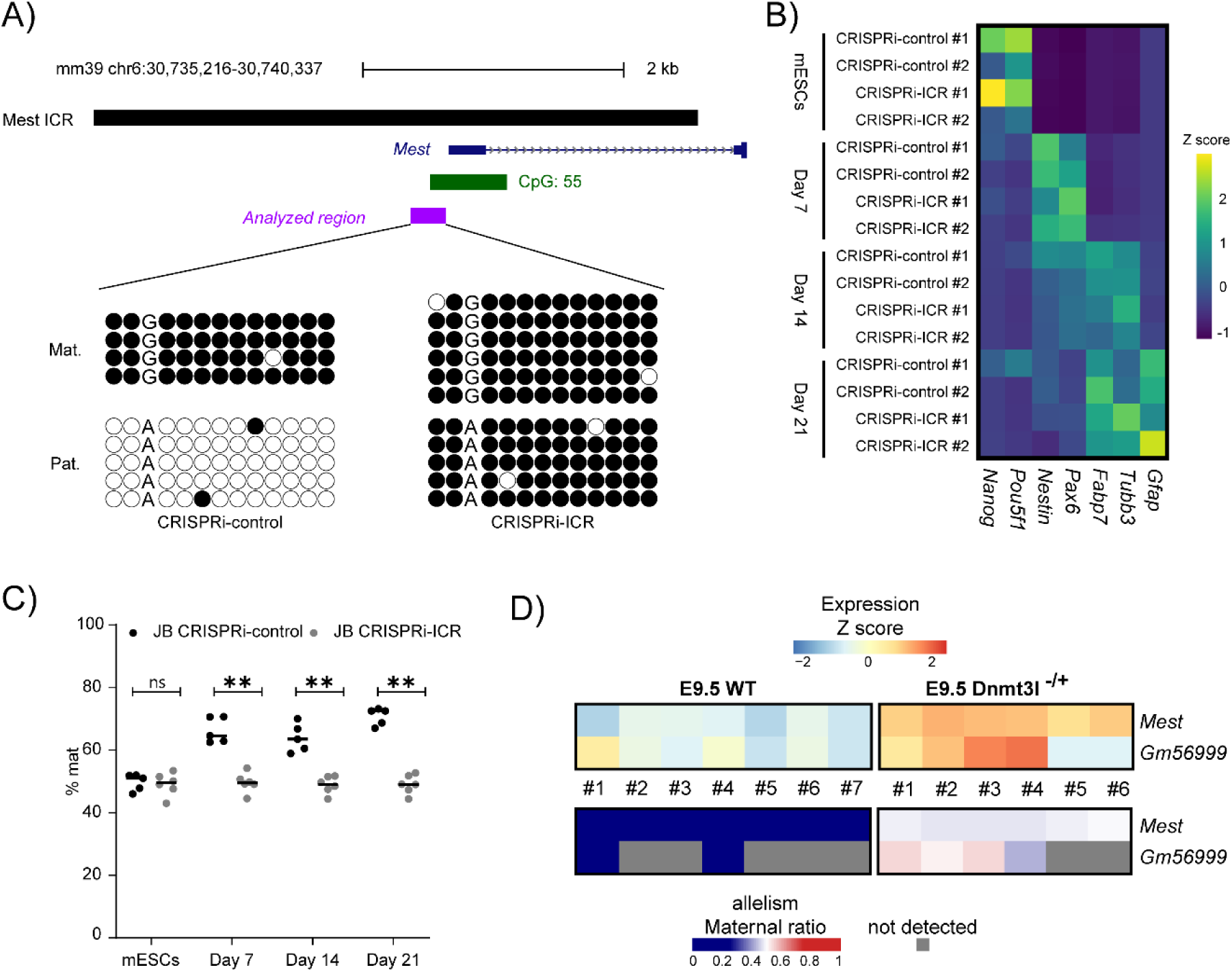
Increased *Copg2* expression in brain organoids derived from CRISPRi-ICR embryonic stem cells. **A)** Schematic of the mouse *Mest* promoter region, illustrating CpG islands (CGI) as annotated in the UCSC Genome Browser. The region analyzed by bisulfite sequencing is marked in purple. Bisulfite sequencing data from CRISPRi-ICR and CRISPRi-control ES cells are shown below, with each row representing CpG dinucleotides on a single chromosome. Filled and open circles indicate methylated and unmethylated CpGs, respectively. Parental origin (Mat., maternal; Pat., paternal) was determined using a strain-specific G/A SNP. **B)** Expression levels of pluripotency and neural markers in CRISPRi-ICR ESCs (n = 4) and at days 7 (n = 3), 14 (n = 3), and 21 (n = 3) of *in vitro* brain organoid differentiation. P-values were determined using an unpaired t-test. **C)** Percentage of *Copg2* maternal expression measured in CRISPRi-ICR and CRISPRi-control J/B embryonic stem cells (ESCs), as well as at days 7, 14, and 21 of *in vitro* brain organoid differentiation. Values were derived from sequencing of RT-PCR products, followed by quantification of ABI format sequence files. Statistical significance was determined with Mann Whitney test (p < 0.05 *, p < 0.01 **, p < 0.001 ***). **D)** Heatmap showing the expression (upper panel) and allelic (lower panel) levels of *Mest* and *MestXL* (assessed through the transcript annotation Gm56999) in E9.5 WT (left panel) and Dnmt3l^-/+^ embryo Data are presented for each independent embryo (WT : n=7 ; Dnmt3l^-/+^ : n=6). Insufficient signal at informative allelic sites was detected in some *Dnmt3L^-/+^* embryos to establish an allelic ratio of Gm56999.

**Supplemental Figure 5:**
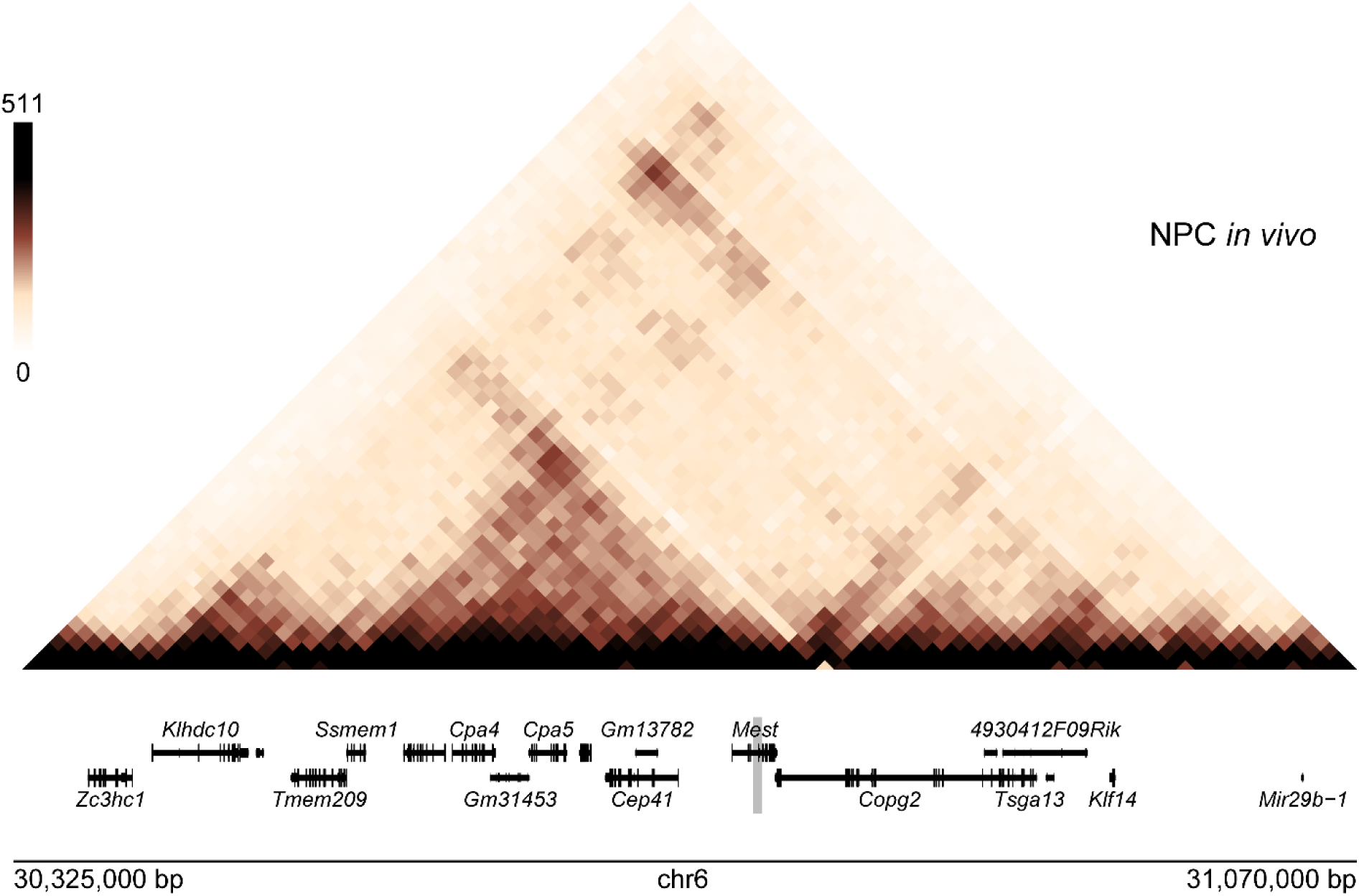
Two chromatin subdomains anchored at the *Mest* ICR. **A)** Genome Browser view of the *Mest/Copg2* domain showing reanalyzed Hi-C data in in vivo NPCs.

**Sup Figure 6:**
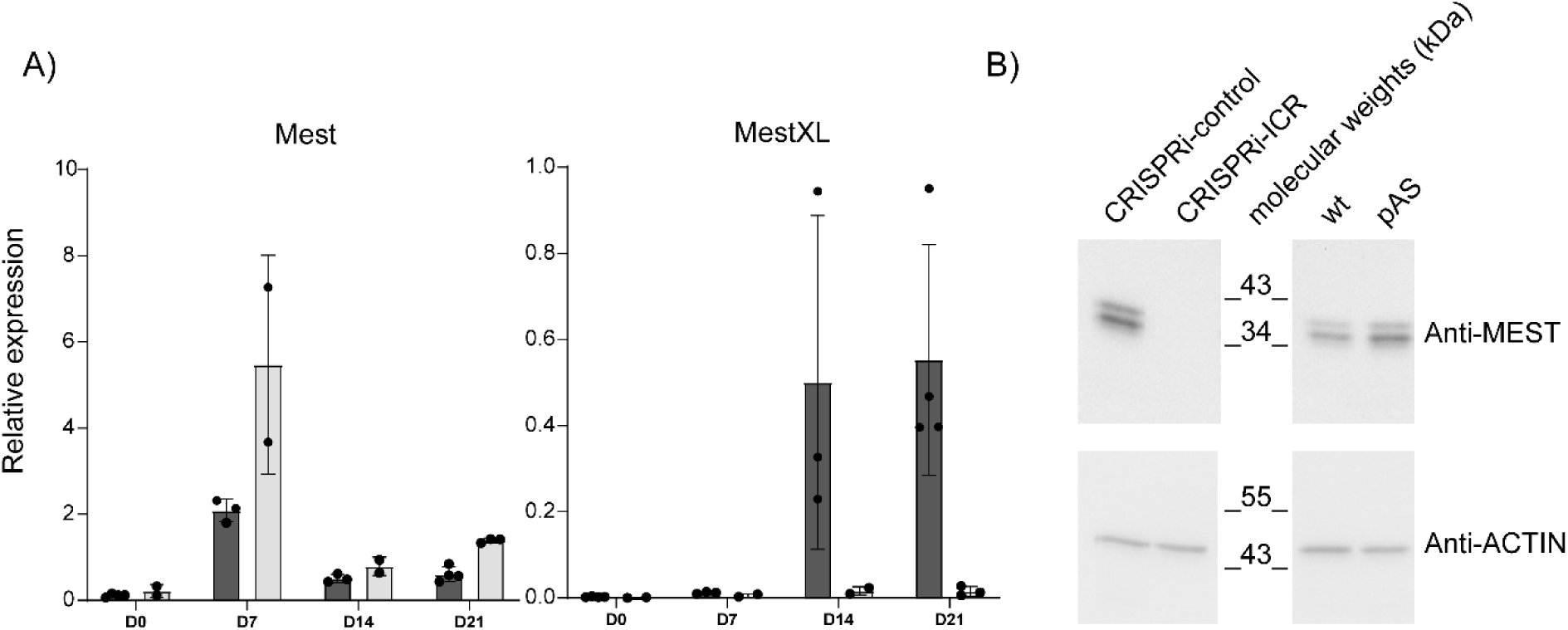
*Mestxl*, but not *Mest,* expression is abolished in pAS cells. A) RT-qPCR analysis of *Mest* and *MestXL* expression levels in ES-pAS cells and at Day 7, Day 14, and Day 21 of in vitro brain organoid differentiation. Values represent the mean of three independent experiments, each analyzed in duplicate. Data are presented as a percentage of expression relative to the geometric mean of three housekeeping genes (*Gapdh*, *Gus*, and *Tbp*), and are shown as mean ± SEM. B) Western blot analysis of *Mest* (upper panel) and β-actin (lower panel) in WT and pAS cells at Day 7. Similar analyses performed in CRISPRi-control and CRISPRi-ICR cells, where *Mest*/MEST expression is abolished, confirm the specificity of the antibody.

